# Biosynthesis of prenylated flavonoids by two membrane-bound prenyltransferases in glandular trichomes of *Macaranga tanarius*

**DOI:** 10.64898/2026.09.23.753730

**Authors:** Ryosuke Munakata, Yoko Maeda, Ryo Shimizu, Akifumi Sugiyama, Shigenori Kumazawa, Shuichi Fukumoto, Kazufumi Yazaki

## Abstract

*Macaranga tanarius* (Euphorbiaceae), a myrmecophilic plant grown in subtropical countries, was identified as a source of Okinawan propolis, a honeybee product, which is glandular trichomes developed on the surface of the fruits. This tissue contains a high amount of characteristic geranylated flavonoids known as nymphaeols (nymphaeol A–C and isonymphaeol B), which exhibit strong antioxidant activity. In this study, we characterized the structure of glandular trichomes of *M. tanarius*, showing that they are sac-like structures containing drupelet-like aggregates of secretary cells, and a subcuticular cavity was also observed. From a cDNA library, we identified two prenyltransferase (PT) genes involved in biosynthesis of nymphaeols, and a PT gene grouped in a primary metabolic PT. Our biochemical experiments demonstrated that one of the former secondary metabolic PTs (MtPT1) is a B-ring-specific geranyltransferase for eriodictyol to yield nymphaeols B and isonymphaeol B. MtPT1 was unusual for a plant PT, an enzyme that yielded multiple reaction products. The other PT (MtPT3) transferred a geranyl moiety to the A-ring of eriodictyol to form nymphaeol A. Although dimethylallyl diphosphate (DMAPP) was not recognized as its prenyl donor substrate of eriodictyol, giving a B-ring geranylated eriodyctiol as the prenyl acceptor substrate, this enzyme used DMAPP to attach the prenyl moiety to the A-ring and produced nymphaeol C. These data suggest that *M. tanarius* has evolved its PT function to produce a diverse range of prenylated flavonoids using a limited set of genes.

## Introduction

Membrane-bound prenyltransferase (PT) plays various important physiological roles in plant primary metabolism, e.g., biosynthesis of plastoquinone and vitamin E necessary for photosynthesis and for antioxidation in seed storage, respectively, and also for the synthesis of ubiquinone, a crucial electron carrier in mitochondria (Munakata and Yazaki, 2024). These enzymes are grouped in the UbiA family, which is responsible for prenylation of aromatic prenyl acceptor substrates. This enzyme family implies that UbiA PTs have divergent substrate specificities, providing a wide variety of plant specialized metabolites. Whereas many different types of phenolics are prenylated, such as coumarins, phloroglucinols, xanthones, and phenylpropanes, a representative group of those metabolites are prenylated flavonoids, in which prenyl residues provide chemical diversities due to the regio-specificity, different chain length, and further modifications. It is also known that prenyl residues often increase the biological activities of flavonoid molecules (Yazaki et al., 2009) .

In biosynthesis of prenylated aromatics, those aromatic substrate PTs take a critical position where shikimate/polyketide pathways couple with mevalonate/MEP (methylerythritol phosphate pathway), major metabolic pathways in plants. Prenylated flavonoids are reported as active substances from many medicinal plants, and chemotaxonomically they mostly occur in Moraceae and Leguminaceae (Fabaceae), while they have been isolated from other plant families as minorities (Yang et al., 2015). Genes coding for those flavonoid-specific PTs have been, therefore, isolated mainly Moraceae and Fabaceae plants (Liu et al., 2025; Sasaki et al., 2008; Yang et al., 2018). In contrast, PT genes responsible for bioactive prenylated flavonoids remain largely unknown in other plant taxa. Recent molecular evolutionary studies of UbiA PTs involved in plant specialized metabolism have demonstrated that the enzymatic function of specialized metabolism-related UbiA-type PTs has diversified in plant taxon-specific manners, and even UbiA PTs with low sequence homology across different plant families catalyze the same reaction, representing an example of convergent evolution (Munakata et al., 2020; Munakata et al., 2021). Thus, the presence of unexplored plant taxa hinders a comprehensive view of the molecular evolution of this enzyme family in the plant kingdom.

*Macaranga tanarius* is a dioecious tropical tree in Euphorbiaceae, which is growing in Okinawa Islands of Japan and other tropical Asian countries (Figure 1A). This plant has been known as a fast-growing tree species and pioneer plant in tropical and subtropical countries. *M. tanarius* is a common tree species in Okinawa and it had been recognized as a non-useful tree for a long time. However, it was identified as the origin of Okinawan propolis, a waxy honeybee product currently used as a food supplement. The source of Okinawan propolis is the glandular trichomes developed on the surface of fruits of this plant that are rich in prenylated flavonoids, nymphaeols (Kumazawa et al., 2008; Kumazawa et al., 2014). The structural features of nymphaeols are that the main flavonoid core is eriodictyol, a flavanone derivative, and all of them have a geranyl residue mostly at the B-ring of the flavonoid skeleton, except for nymphaeol A (Figure 2). Nymphaeol C has a dimethylallyl residue at the A-ring in addition to a geranyl residue at the B-ring. This plant species is expected to be a valuable source of bioactive prenylated flavonoids, while so far no aromatic PT genes have been isolated in Euphorbiaceae.

**Figure 1.**
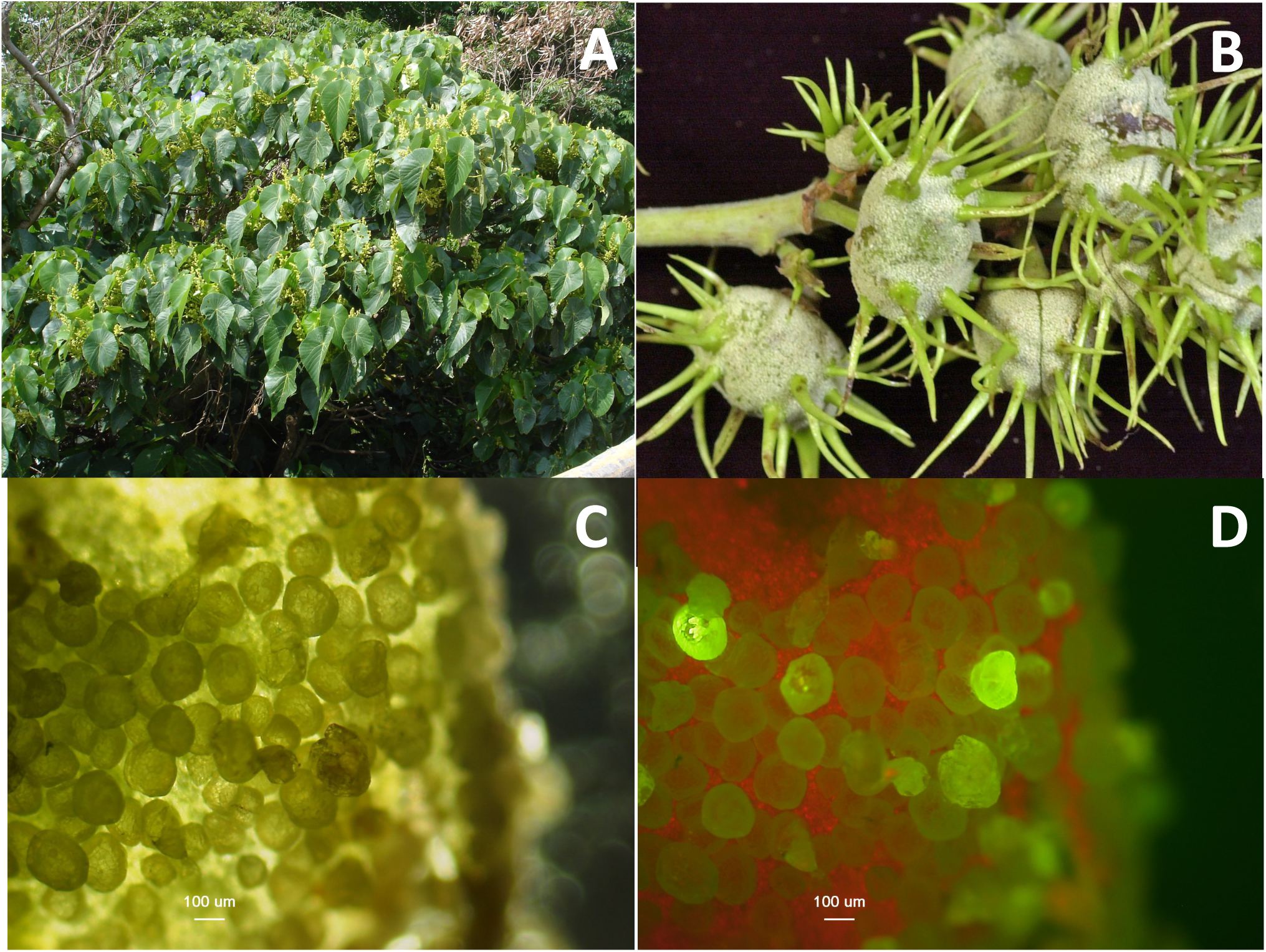
*Macaranga tanarius* grown in Okinawa Island. (A) A male tree of *M. tanarius*. (B) Fruits of *M. tanarius* covered by white glandular trichomes where a trace of scratching by honeybees is also seen. (C) Enlarged image of the fruit surface. (D) Fruit surface under UV irradiation.

**Figure 2.**
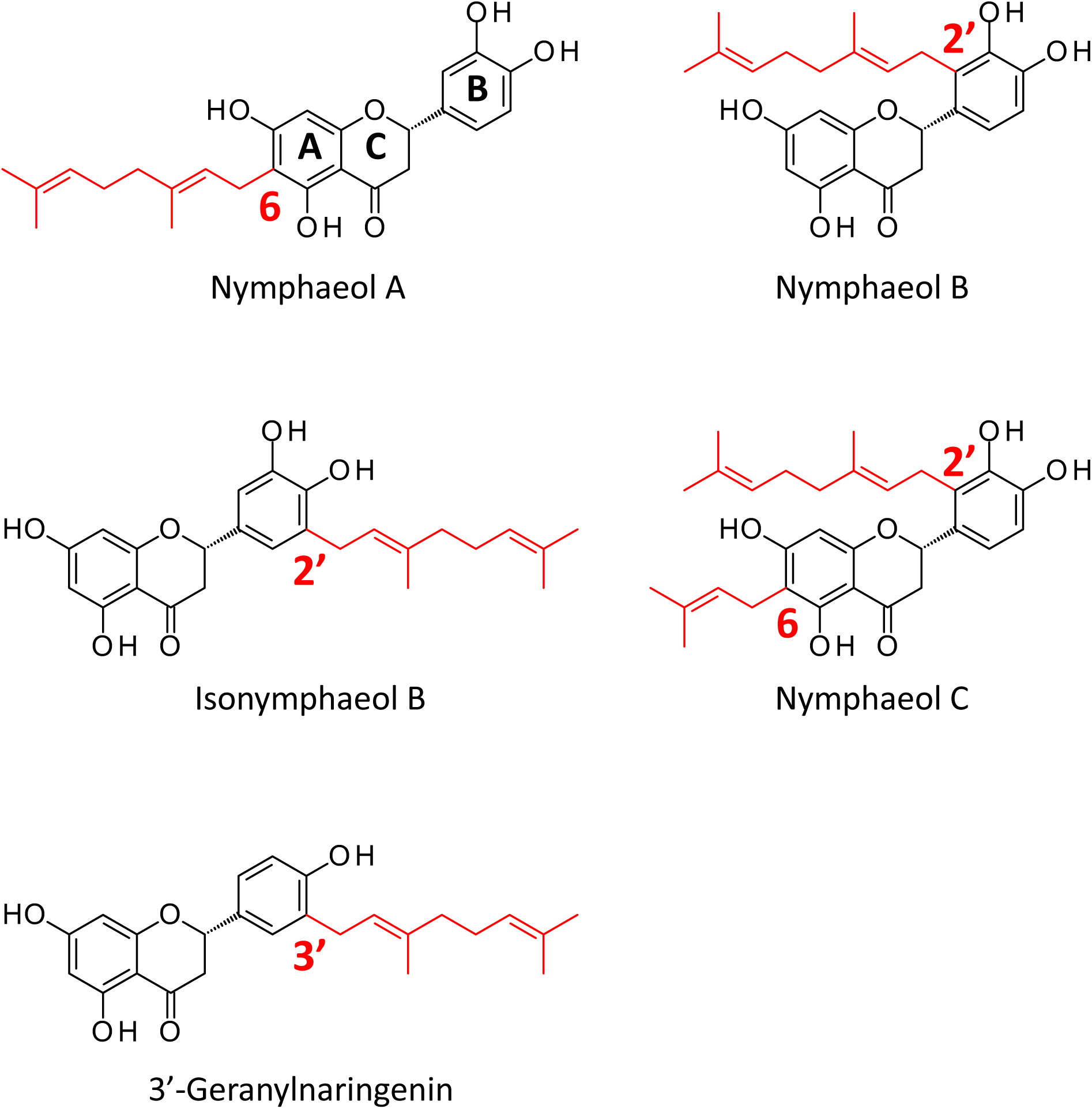
Prenylated flavonoids commonly present in *M. tanarius* and Okinawan propolis. Prenyl moieties were highlighted in red.

In the present study, we microscopically characterized glandular trichomes of *M. tanarius* and prepared a glandular trichome cDNA library. Two UbiA-type PT genes isolated based on the library have been expressed in *Nicotiana benthamiana* by a transient expression system, and characterized their enzymatic functions, as well as their subcellular localization. Our study provides new insights into the UbiA PT family involved in plant specialized metabolism.

## Materials and Methods

### Reagents

Dimethylallyl pyrophosphate (DMAPP) and geranyl pyrophosphate (GPP) used in the preliminary trials were kindly provided by Dr. Hirobumi Yamamoto (Toyo University), and Dr. Tomohisa Kuzuyama (The University of Tokyo) and Dr. Takashi Kawasaki (Kyoto University), respectively. Prenyl diphosphates used in the other assays were purchased from Sigma-Aldrich (St. Louis, MO, US). The aromatic molecules were purchased from Cayman Chemical (Ann Arbor, Michigan, US), Combi-Blocks (San Diego, CA, US), Extrasynthase (Lyon, France), FUJIFILM Wako Pure Chemicals (Osaka, Japan), Sigma-Aldrich, and Tokyo Chemical Industry (Tokyo, Japan). [1-^14^C] DMAPP was purchased from American Radiolabeled Chemicals, Inc. Authentic standards of nymphaeols were prepared as previously published (Kumazawa et al. 2008).

### Structures of glandular trichomes of *M. tanarius*

The fruits of *M. tanarius* were harvested on 23^rd^ July, 2008, on Okinawa Island. The surface was sliced with a scalpel and observed with a fluorescent microscope Axoiskope 2 (Zeiss, Jena, Germany). Bright filed pictures were taken without a filter (Figure 1C), and for fluorescence observation, a long-pass filter for green fluorescent protein (exciation: BP 470/20 nm, beamsplitter: FT 493nm, BP500-530 nm) was used (Figure 1D). Observation of glandular trichomes using a Variable Pressure-Scanning Electron Microscope (VP-SEM) was performed as follows. Fruit surface was cut out (approximately 5 mm square) and the fruit peel sample was affixed to the sample holder using wood glue. Glandular trichomes were observed using VP-SEM (S-3500N with cool stage, Hitachi, Japan) without pre-treatment (Figure 3A and B). When observing the underside of the glandular scales, we used adhesive tape to peel the scales off the fruit surface and secured them, along with the tape, to the stage for observation (Figure 3C and D). The abaxial side of the leaf was also observed in a similar manner (Figure 3E and F). Resin embedding and section preparation were performed as follows. The surface of fresh fruits was cut and soaked in 30 % ethanol in a PCR tube and stand for 1 h at room temperature. Then ethanol concentration was increased stepwise (50, 70, 90, 100 %), with each treatment lasting 1 h. Then, the ethanol was substituted with 50 % Technovit 7100 (Heraeus Kulzer GmbH, Wehrheim, Germany) in ethanol and soaked overnight. The sample was then placed in 5 ml Technovit 7100 containing 0.05 g hardener I and stand for 8 h at room temperature. Then, the solution was substituted with Technovit 7100 (7.5 ml) containing hardener II (0.5 ml) to solidify overnight. The resin containing sample was affixed to a histology block (Heraeus Kulzer GmbH) using Technovit 3040 (Kulzer). The sampling surface was trimmed with a dental drill (viva-mate3, Nakanishi, Japan), and sliced to a thickness of 8 µm thickness using a microtome (LEICA RM2155). The samples were observed with Axioscope 2 as described above.

**Figure 3.**
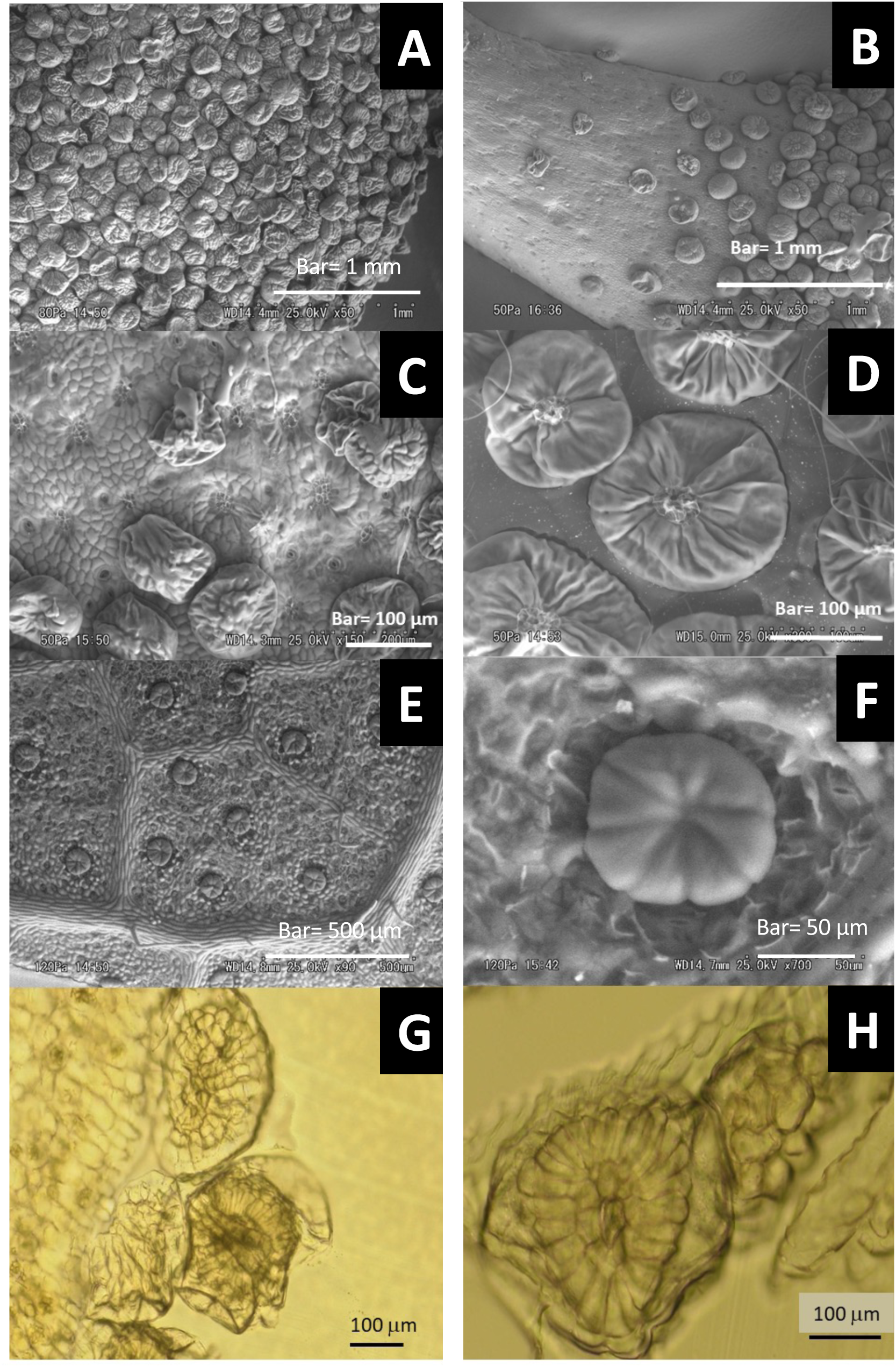
Micrographs of glandular trichomes of *M. tanarius*. (A) Fruit surface of glandular trichomes. (B) A soft thorn of a fruit. (C) Epidermal cells and traces of stalk cells of glandular trichomes of a fruit. (D) An upside-down image of fruit glandular trichomes. (E and F) Glandular trichomes of a leaf (abaxial side). (G and H) Cross-sections of resin-embedded glandular trichomes from a fruit.

### Native enzyme assay

Fresh plant materials (male flowers and leaves) were frozen with liquid nitrogen and crushed to fine powder with a mortar and pestle. The powder was homogenized in a 100 mM K-Pi buffer (pH 6.4, 30 ml) containing 1 mM dithiothreitol, polyvinylpolypyrrolidone (0.1 g g^-1^ fresh sample), followed by centrifugation at 3,000 × *g* for 5 min at 4 °C to obtain the supernatant. After filtration with Miracloth (Merck) the filtrate was further centrifuged at 8,000 *g* for 20 min at 4 °C to obtain the pellet and the supernatant (soluble fraction). The pellet was suspended in a 100 mM Tris-HCl buffer (pH 7.5, 300 µl) containing 1 mM dithiothreitol and 1 mM phenylmethylsulfonyl fluoride to yield the organellar fraction. Enzyme reaction was done with [1-^14^C] DMAPP (55 mCi mmol^-1^) with each flavonoid substrate (1.0 mM) in the presence of 10 mM MgCl2 at 30 °C for 30 h. The reaction was terminated by the addition of 3 M HCl and extracted with 1 ml ethyl acetate. The organic phase was evaporated, and 30 µl methanol was added to dissolve the residue. The entire amount was spotted onto a TLC plate (20 × 20 cm, Kiesel gel 60F254, MERCK), which was then developed with a solvent system (toluene : ethyl acetate : acetic acid = 70 : 30 : 0.3). After drying the TLC plate was set with an imaging plate in BAS1800 for autoradiography. After 5 days, images were analyzed.

### cDNA library construction and isolation of PT cDNAs

RNA sample was prepared from glandular trichomes of *M. tanarius*, which were collected by scratching the surface of fresh fruits. As commercially available RNA extraction kits did not work as expected, we employed a conventional cetyltrimethylammonium bromide (CTAB) method, and 1.35 µg total RNA was used to construct a cDNA library. Because of the limited amount of RNA sample, we used SMART (Switching Mechanism At 5′End of RNA Template) method for the plasmid library construction according to the manufacturer’s instruction (Takara Bio, Japan). Because our target was PT genes encoding membrane-bound proteins, we used the yeast-*Escherichia coli* shuttle vector pDR196 (Sasaki et al., 2008), to which the SfiIA and SfiIB sites were introduced between the EcoRI and XhoI sites. The plasmids were introduced into *E. coli* DH10B strain by electroporation, and amplification was performed using this strain. The titar was 1.3 × 10^10^ cfu ml^-1^.

Sequencing was outsourced to Takara-Bio (Kusatsu, Japan). Out of total read 10,080, filtering of nucleotide sequence data obtained through base calling by Phred using the PFP (paracel filtering package, Paracel, Vietnam) software resulted in 9,858 reads that have sequence length longer than 300 bp. Clustering analysis was done to yield the results, 1,366 clusters composed of 6,465 reads and 3,393 singlets (Supplementary Figure S2). In total, 4,759 non-redundant EST sequences were generated.

To find out candidate genes encoding flavonoid-substrate PTs, blastx was done using homogentisate PT, which is involved in vitamin E biosynthesis and an ancestral gene for secondary metabolic PTs, as a query. Sequences lacking conserved domains important for catalytic function, i.e. aspartate-rich motifs, were eliminated from the candidate despite an appreciable sequence similarity (Munakata and Yazaki 2024). Three genes remained as candidates: *Macaranga tanarius PT1* to *3* (*MtPT1–3*).

### *In silico* analysis of polypeptides

Amino acid identities among PTs were calculated based on ClustalW multiple alignments using BioEdit. Transit peptides (TPs) and transmembrane regions were predicted with DeepLoc2.1 (https://services.healthtech.dtu.dk/services/DeepLoc-2.1/) ((Ødum et al., 2024) and DeepTMHMM (https://dtu.biolib.com/DeepTMHMM/) (Hallgren et al., 2022), respectively. PT polypeptide sequences were collected based on our previous work (Matsushita et al., 2026) and aligned for phylogenetic analysis using MAFFT version 7 (https://mafft.cbrc.jp/alignment/software/) (Kuraku et al., 2013; Katoh et al., 2019). A maximum-likelihood phylogenetic tree was constructed with IQtree version 1.6.11. (https://iqtree.github.io/) (Nguyen et al., 2015), using the model JTT+F+R6 chosen by ModelFinder (Kalyaanamoorthy et al., 2017). The Shimodaira–Hasegawa approximate likelihood ratio test (SH-aLRT test, 1,000 replicates) (Guindon et al., 2010) and the ultrafast bootstrap test (UFBoot, 1,000 replicates) (Minh et al., 2013) were performed to evaluate branch support.

### Transient expression of GFP-fusion proteins and microscopic observation

Nucleotide sequences corresponding to the N-terminal regions (66 amino acids of MtPT1, designated MtPT1TP; 75 amino acids of MtPT3, designated MtPT3TP) were amplified by PCR using KOD-plus (Toyobo, Osaka, Japan). Primer pairs for *MtPT1TP* (Fw, 5′-CACCATGGTTGCATCAAGCA -3′; Rv, 5′-CAACCCATCTTC GCTGCTAT-3′) and MtPT3TP (Fw, 5′-CACCATGTTTTTTCAATCTT-3′; Rv, 5′-AGAATTTGGATGTGCATTTT-3′) were used for PCR amplification. The resulting DNA fragments were then subcloned into the pENTR/D-TOPO vector (Thermo Fisher Scientific, Waltham, MA, US). Subsequently, these sequences were introduced into the destination vector pGWB505 to fuse with green fluorescent protein (GFP) (Nakagawa et al., 2007) by LR recombination to yield *CaMV35Spro-MtPT1TP-sGFP* and *CaMV35Spro-MtPT3TP-sGFP*.

Agroinfiltration was performed as previously described (Karamat et al., 2014), and transiently expressed MtPT1TP-sGFP and MtPT3TP-sGFP were analyzed using fluorescence microscopy. Microscopic analysis was performed essentially as described (Matsushita et al., 2026). The plasmid pHKN29 (Kumagai and Kouchi, 2003) containing *CaMV35Spro-GFP* was used as a control. For each construct, multiple cells showing GFP signals derived from multiple leaves were observed, and all cells gave the same results.

### Construction of plant expression plasmids for enzymatic characterization

The coding sequences (CDSs) of *MtPT1* and *MtPT3* were amplified using KOD-FX-neo and KOD-plus (Toyobo), respectively, and the primer pairs for *MtPT1* (Fw, 5′-CACTGTTGATACATATGGTTGCATCAAGCATGCT-3′; Rv, 5′-ATTCAGAATTGTCGATCAATTCATAAATATAGATA-3′) and *MtPT3* (Fw, 5′-CACTGTTGATACATATGTTTTTTCAATCTTCTTCGC-3′; Rv, 5′-ATTCAGAATTGTCGATCAGTTCATAAATATAGGAAGC-3′). The amplicons were subjected to in-fusion reactions (Takara-bio, Kusatsu, Japan) together with the pRI201-AN (Takara-bio) that was linearized by double digestion with SacI and NdeI, yielding the plasmids containing *CaMV35Spro-Arabidopsis thaliana alcohol dehydrogenase (AtADH) 5’-UTR-MtPT1* and *CaMV35Spro-AtADH 5’-UTR-MtPT3*.

### Transient expression of *MtPTs* in *N. benthamiana* leaves and microsome preparation

pRI201-AN-*MtPT1*, pRI201-AN-*MtPT3*, and pRI201-AN-empty vector (negative control) were introduced into *Agrobacterium tumefaciens* strain LBA4404. The transformants were infiltrated into *N. benthamiana* leaves together with *A. tumefaciens* strain C58C1 harboring pBIN61-P19 (Norkunas et al. 2018). Microsomes were prepared essentially as described (Karamat et al., 2014) and finally suspended into the reaction buffer (100 mM Tris-HCl (pH 7.5) containing 1 mM dithiothreitol) and stored at -80 °C until use.

### *In vitro* assay conditions and extraction of reaction products

The frozen microsomes were thawed under running water and diluted with the reaction buffer. The standard reaction mixture (200 µl) was composed of 200 µM prenyl acceptor substrate, 200 µM prenyl donor substrate, 10 mM MgCl2, and microsomes containing MtPT1 or MtPT3 (ca. 5-µg protein calculated by Bradford assay) (Bradford, 1976). The reaction mixture was incubated at 28 °C overnight. All *in vitro* experiments were performed at least twice to confirm their reproducibility. Substrate specificity tests were performed in three independent incubations. Sequential prenylation of eriodictyol to yield nymphaeol C was carried out by co-adding microsomes containing MtPT1 and MtPT3 in the reaction tube.

The reaction was terminated by the addition of 100 µl of 3 M HCl. Aromatic molecules were extracted using 300 µl of ethyl acetate. The mixture was vortexed for 5 min and centrifuged for 5 min to separate the water and the ethyl acetate phases. The upper ethyl acetate phase (200 µl) was collected and dried by vacuum evaporation. After dissolving in 100 µl of methanol, the extract was subjected to liquid chromatography-photodiode array (LC-PDA) and LC-tandem mass spectrometry (LC-MS^2^) analyses.

### LC/PDA and LC/MS^2^ analyses of reaction products

The extracts of incubation mixtures were analyzed with LC-PDA (NEXERA UHPLC system, Shimadzu, Kyoto, Japan) and LC-MS^2^ (Vanquish UHPLC System coupled to a Q Exactive Focus mass spectrometer, Thermo Fisher Scientific) as described (Matsushita et al., 2026), except for the use of a C18 reverse-phase column (ACQUITY UPLC BEH C18 1.7 μm, 2.1 × 150 mm, Waters) connected to a guard column (ACQUITY UPLC BEH C18 VanGuard Pre-column (1.7 μm, 2.1 × 5 mm, Waters) in LC/MS^2^ analysis in this study. In LC/MS^2^ analysis of the reaction mixtures of the eriodictyol geranyltransferase assays, the reaction products, except for one unidentified product from MtPT1, were identified by direct comparison with nymphaeol standards. The other reaction products were searched based on their molecular ion peaks in MS spectra, as well as their fragmentation relevant to the prenyl moiety attached to the aromatic ring in the MS^2^ spectra. These fragmentations were characterized by the neutral losses of 56, 124, 136, and 192 daltons, corresponding to a *C*-dimethylallyl moiety, a *C*-geranyl moiety, an *O*-geranyl moiety, and a *C*-farnesyl moiety, respectively (Simons et al., 2009; Munakata et al., 2014; Munakata et al., 2021).

Quantification of reaction products was carried out based on areas close to λ max (genistein, 260 nm; flavanones and taxifolin, 290 nm; umbelliferone, 320 nm; luteolin, 340 nm; butein, 380 nm) in LC/PDA analysis. Due to the instability of nymphaeols and the lack of standards for the other products, all reaction products were quantified as substrate equivalents. If background signals were present in the negative control, they were subtracted from the calculated activities. In substrate specificity analysis, enzymatic activities less than 5 % compared to the highest one were considered trace activities.

## Results

### Structures of glandular trichomes of *M. tanarius*

Prenylated flavonoids in fruits are exclusively accumulated in the glandular trichomes (Kumazawa et al., 2014), which densely cover the surface of fruits (Figure 1B and C), where chloroplast-derived red fluorescence was not observed under ultraviolet light (Figure 1D). Autofluorescence of chlorophyll is observed in epidermal cells of fruits, whereas glandular trichomes exhibit green fluorescence instead of chlorophyll-derived red fluorescence. The characters of prenylated flavonoids in *M. tanarius*, known as nymphaeols, are that most prenyl residue attached on the aromatic ring is geranyl and many of them are found in the B-ring of the flavonoid. The main flavonoid core of nymphaeols is eryodictiol, and as a minority prenylated naringenin is also found (Figure 2).

The surface of glandular trichomes was observed with a VP-SEM (Figure 3). Glandular trichomes of *M. tanarius* show uniform shape, i.e. slightly flattened round shape of 160 ± 20 µm (n = 20) in diameter (Figure 3A–D). They developed densely on the surface of the entire fruit but not on the soft thorns (Figure 1A and B). To observe the stalk cells, glandular trichomes were stripped off with a sticker (see Materials & Methods) and observed under VP-SEM, which showed several cells gathering at one locus by which the sack-shaped glandular trichomes were anchored to the fruit surface (Figure 3C and D). The trace of these stalk cells was also seen (Figure 3C). Glandular trichomes were also developed on the abaxial side of the leaves at a lower density than on the fruits (Figure 3E and F), while the size and shape were very similar to those developed on the fruits (Figure 3G).

Fresh glandular trichomes are then embedded into a resin and sections are made by microtome to observe the cell organization inside (Figure 3G and H). In the glandular trichomes a unique cell organization was observed, i.e., a cluster composed of multiple cells with radial structures was formed, and an appreciable space outside the clustered cells under the cuticle was observed. This structure is different from glandular trichomes of hops and spearmint, in which secretary cells are aligned at the bottom of the inner space of the glandular trichome (Ramos et al. 2024), but the cell organization within the glandular trichomes is often plant species-specific.

### Detection of PT activity

Cell-free extract was prepared from flowers and leaves of *M. tanarius*, which was fractionated into organelle-rich and soluble fractions (supernatant of 100,000 × *g*), as well as microsomes. Enzymatic reaction was done with ^14^C-DMAPP and flavonoids (eryodictiol or 2′-geranyl-eryodictiol) as the prenyl donor and acceptor substrates, respectively. After the reaction, the products were analyzed by autoradiography using normal phase TLC plates. Among tested crude enzyme samples, only organelle-rich fraction prepared both from flowers and leaves gave hydrophobic products, suggesting the existence of PTs in organelles (Supplementary Figure S1). However, the enzyme activity was weak and the yield of protein from leaves was very low. It is also not feasible to collect a reasonable amount of glandular trichomes from the fruit surface for biochemical studies. Thus, further biochemical analysis was not performed, and we shifted to transcriptomic screening.

### Isolation of PT cDNAs from a library prepared from glandular trichomes

To isolate candidate PT genes involved in biosynthesis of nymphaeol derivatives, we constructed a cDNA library from glandular trichomes where these molecules are exclusively accumulated. We collected glandular trichomes from fresh fruits and prepared total RNA. Although several widely utilized methods were applied to isolate total RNA, no substantial amount of RNA was collected. Thus, we applied a conventional CTAB method, resulting in a yield of 12.1 µg of total RNA from approximately 4 g of glandular trichomes of the fruit surface.

Due to the limited amount of RNA sample, we prepared a cDNA library with the SMART method, in which full-length cDNAs are likely condensed (see Materials and Methods). In the construction of the cDNA library, we employed the yeast-*E. coli* shuttle vector pDNR 196, because the candidate cDNAs encode membrane-bound proteins. Out of the total read 10,080, 3,393 singlet clones and 1,366 clusters were obtained after filtering and clustering (Supplementary Figure S2).

PTs that accept aromatic substrates in plant specialized metabolism are classified into several groups based on their possible ancestral sequences in primary metabolism: mitochondria-localized *p*-hydroxybenzoate polyprenyltransferase involved in ubiquinone biosynthesis (Ohara et al., 2006), plastid-localized homogentisate phytyltransferase involved in vitamin E biosynthesis (Sattler et al., 2004), and plastid-localized homogentisate solanesyltransferase involved in plastoquinone biosynthesis (Norris et al., 1995). All family members possess a couple of common features, i.e., aspartate-rich motif, such as NDxxDxxxD or NQxxDxxxD, which are responsible for binding to the prenyl donor substrate. However, these three classes have low amino acid identities (approximately less than 30 %). Taking the advantage of our previous knowledge and sequence collections encoding PTs for aromatic substrates, we narrowed down candidate clones that code for PTs for flavonoids substrates. As a result, we obtained cDNA fragments for five flavonoid PT candidates from the cDNA library. Out of them, we cloned three cDNAs containing full coding sequences by rapid amplification of cDNA ends (RACE), which were designated *MtPT1–3*.

MtPT1-3 show approximately 27 % (MtPT1 vs 2), 64 % (MtPT1 vs 3), and 31 % (MtPT2 vs 3) amino acid identities each other. Further i*n silico* analyses of these polypeptides predicted that MtPT1–3 possess multiple transmembrane helices and a transit peptide (TP) in their N-terminal regions (Supplementary Figure S3), which are typical features among plant membrane-bound PTs. In the phylogenetic relationship among plant aromatic-substrate PTs, MtPT1 and MtPT3 appeared to form an independent group (Figure 4). Berberidaceae PTs are relatively close to MtPT1 and 3, but their amino acid identities are not high (30–32 %), suggesting that MtPT1 and MtPT3 show enzymatic functions responsible for specialized metabolism of *M. tanarius*. In contrast, MtPT2 was grouped in the clade of VTE2-1, which is homogentisate phytyltransferase responsible for vitamin E formation in primary metabolism (Sattler et al., 2004). Therefore, we characterized MtPT1 and MtPT3 for further analysis.

**Figure 4.**
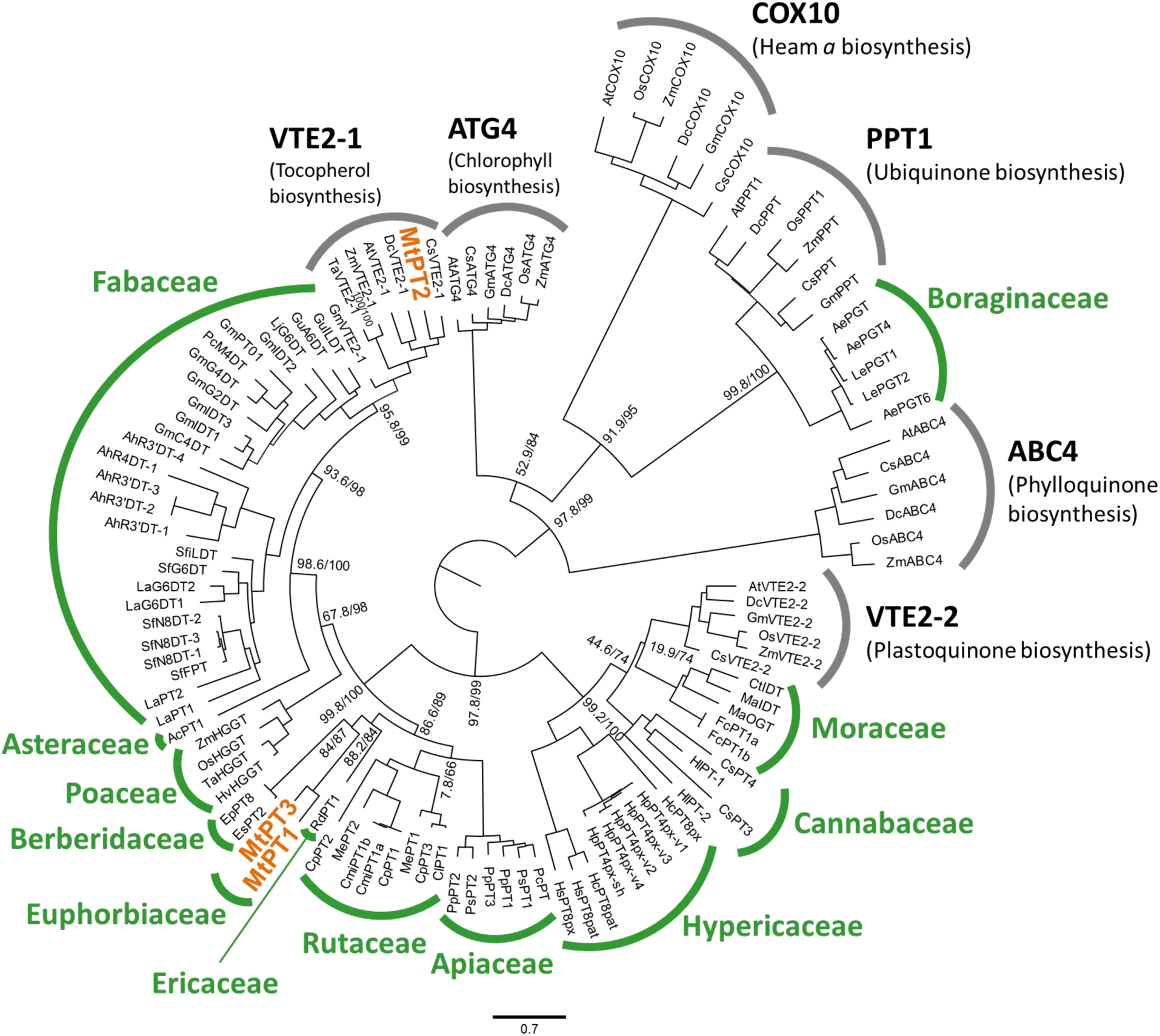
Phylogenetic relationship of MtPT1–3 in the UbiA PT family. A maximum likelihood phylogenetic tree was constructed based on a MAFFT multiple alignment of plant UbiA PT polypeptide sequences. The results of the SH-aLRT test (1,000 replicates, left) and the UFBoot test (1,000 replicates, right) are shown at the nodes separating the clades. Clades containing PTs involved in primary metabolism and specialized metabolism are highlighted in gray and green, respectively. The scale bar indicates an amino acid substitution rate per site of 0.7. The plant origins and the accession numbers of the PT sequences are listed in our previous work (Matsushita et al., 2026).

### Subcellular localization of MtPT1 and MtPT3

To experimentally assess the subcellular localization of MtPT1 and MtPT3, the N-terminal regions of MtPT1 (66 aa.) and MtPT3 (75 aa.), which possibly contain their TPs, were fused to synthetic GFP (MtPT1TP-sGFP and MtPT3TP-sGFP). These GFP-fusion proteins were transiently expressed in *N. benthamiana* leaves by agroinfiltration. Confocal microscopic analysis of epidermal cells of infected leaves indicated that both MtPT1TP-sGFP and MtPT3TP-sGFP were localized in chloroplasts, depending on the MtPT1TP and MtPT3TP sequences (Figure 5). This result suggests that both MtPT1 and MtPT3 function in plastids of *M. tanarius*, where DMAPP and GPP are provided via the methylerythritol phosphate pathway.

**Figure 5.**
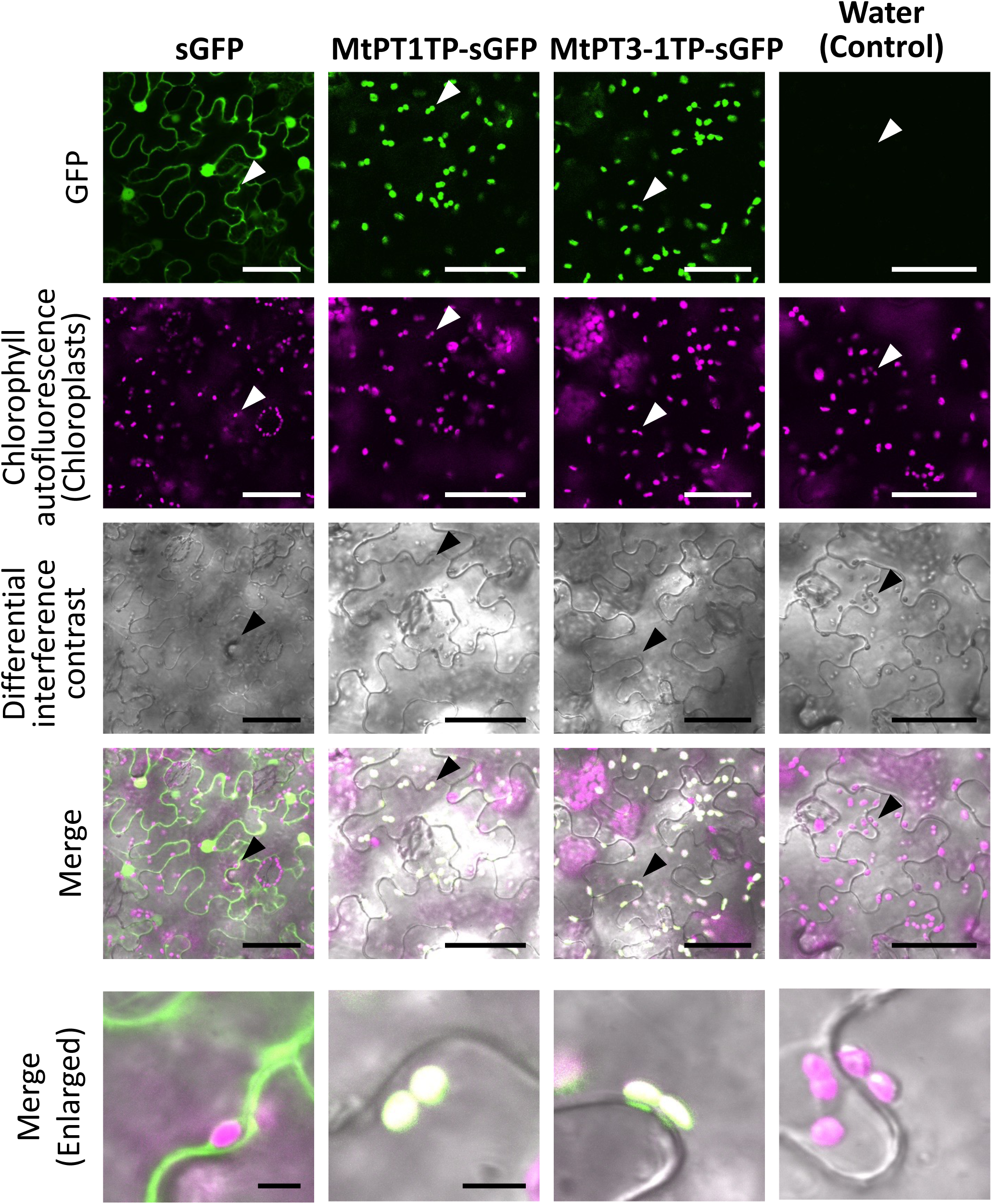
Subcellular localization of MtPT1 and MtPT3. MtPT1TP-GFP and MtPT3TP-sGFP, free GFP as a control were transiently produced in *N. benthamiana* leaves by agroinfiltration. Water infiltration was performed as another control. The localization of the GFP signal was observed using confocal microscopy, and chloroplasts were visualized based on chlorophyll autofluorescence shown in magenta (pseudocolor). The brightness and contrast were adjusted in an unbiased manner among all samples. The regions of interest indicated by the arrowheads were enlarged and are shown in the bottom row. The scale bars indicate 50 µm and 5 µm (enlarged images).

### Transient expression of MtPT1 and MtPT3 and detection of their enzymatic functions

Although we initially considered yeast as an expression host of PTs, it has been reported that microbial expression systems often cause problems in the functional expression of membrane-bound PTs (Karamat et al., 2014; Yang et al., 2018). Thus, we selected *N. benthamiana* as a host organism to express the full CDSs of *MtPTs*. *MtPT1* and *MtPT3* were subcloned into the binary vector pRI201-AN for the expression under the control of the *CaMV35Spro* and *AtADH* 5’-UTR as a translation enhancer. These plasmids were introduced into *N. benthamiana* leaves via agroinfiltration. An *Agrobacterium* strain harboring pBIN61-*P19*, a suppressor of gene silencing, was co-infiltrated to boost the expression of *MtPTs* (Norkunas et al., 2018). Four days after infiltration, infected leaves were homogenized and ultrancentrifuged to obtain microsomal fractions, which were used as crude enzymes in the PT assays.

As a native substrate occurring in *M, tanarius*, eriodictyol was used as a prenyl acceptor, and GPP was used as a prenyl donor substrate because geranylated forms of this flavonoid molecule are the vast majority in glandular trichomes of *M. tanarius* (Kumazawa et al., 2008). When MtPT1 was incubated with this substrate pair in the presence of MgCl2 as a cofactor, multiple products (1ErioG-P1–4) were yielded with 1ErioG-P1 and P2 being the major products. 1ErioG-P1, P2, and P3 were identified as nymphaeol B, isonymphaeol B, and nymphaeol A, respectively, by direct comparison of their retention times and MS and MS^2^ spectra with those of standard specimens (Figure 6 and Supplementary Figure S4A–F), demonstrating that MtPT1 preferentially transfers prenyl chains to the B-ring. The minor product 1ErioG-P4 was predicted as an *O*-geranylated eriodictyol, but we were unable to identify its chemical structure due to lack of the corresponding standard (Supplementary Figure S4G and H).

**Figure 6.**
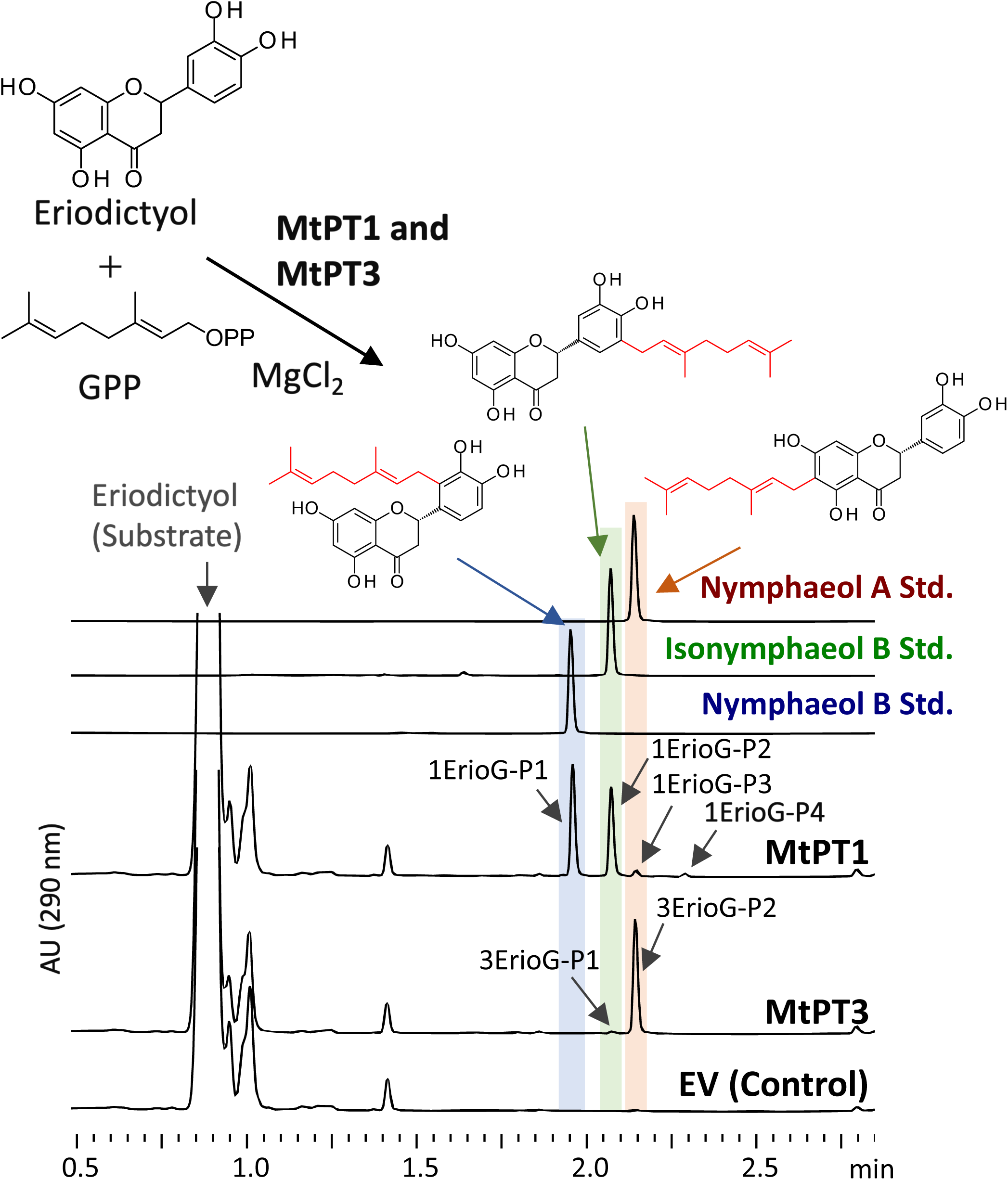
Eriodictyol geranyltransferase activities of MtPT1 and MtPT3. Ultraviolet chromatograms (290 nm) of the reaction mixtures of recombinant MtPT1 and MtPT3 in use of eriodictyol and GPP as a substrate pair. MtPT1 and MtPT3 yielded four (1EripG-P1–4) and two (3ErioG-P1 and P2) products, respectively. Empty vector was used as a control.

Under the same reaction conditions, MtPT3 produced two enzymatic reaction products, 3ErioG-P1 and P2. The minor product 3ErioG-P1 and the major product 3ErioG-P2 were identified as isonynphaeol B and nymphaeol A, respectively, as in the case of MtPT1 (Figure 6 and Supplementary Figure S4C–F). These results indicated that MtPT1 and MtPT3 both show multiple geranyltransferase activities for eriodictyol with different regio-specificities each other, *i.e.*, MtPT1 and MtPT3 predominantly geranylate the B-ring and the A-ring of eriodictyol, respectively.

### Substrate specificity of MtPT1 and MtPT3

Substrate specificities of MtPT1 and MtPT3 were assessed using different aromatic molecules in the presence of GPP as a prenyl donor (Figure 7). Among the flavanone molecules, MtPT1 accepted naringenin, the eriodictyol analog without the hydroxyl moiety at the 3′ position, with a high efficiency (Supplementary Figure S5A). MS^2^ spectra showed the major signal at *m/z* = 153, which is derived from the C-ring-cleaved molecule possessing a non-prenylated A-ring (Supplementary Figure S5B)(Ameer et al., 1996). Another characteristic signal is *m/z* = 285, generated by the neutral loss of 124 daltons, and this fragmentation is specific to a *C*-geranyl moiety attached to the aromatic ring (Simons et al., 2009; Munakata et al., 2014). Moreover, MtPT1 showed a high specificity to the B-ring for eriodictyol. In principal, UbiA PTs catalyzing aromatic *C*-prenylation transfer a prenyl moiety to a carbon atom ortho to a phenolic hydroxyl moiety through Friedel-Crafts alkylation (Leveson-Gower and Roelfes, 2022). Based on this evidence, the MtPT1 product is assumed to be 3′-geranylnarignenin (Supplementary Figure S5C), a chemical component of both *M. tanarius* glandular trichomes and Okinawan propolis (Kumazawa et al., 2008).

**Figure 7.**
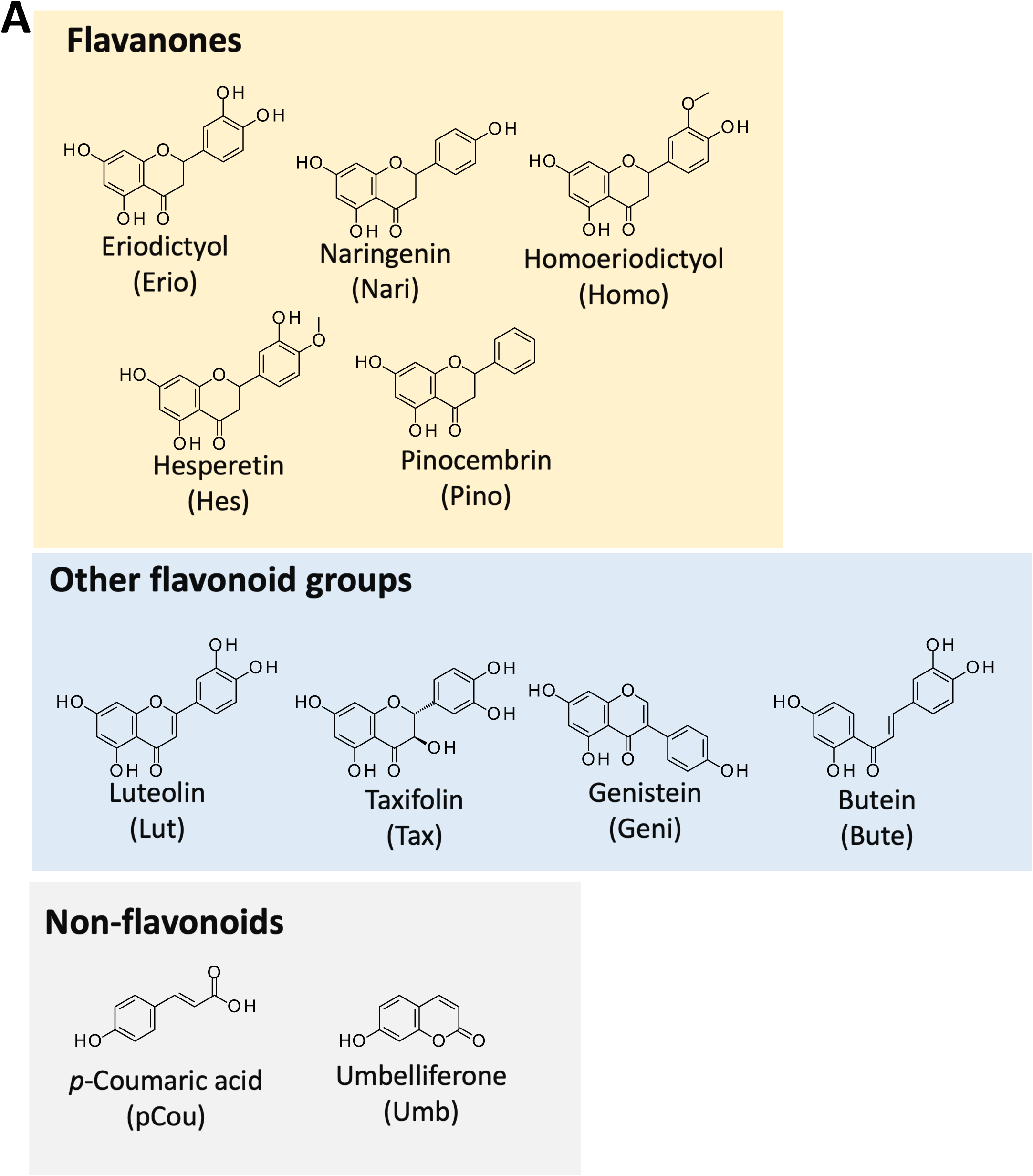
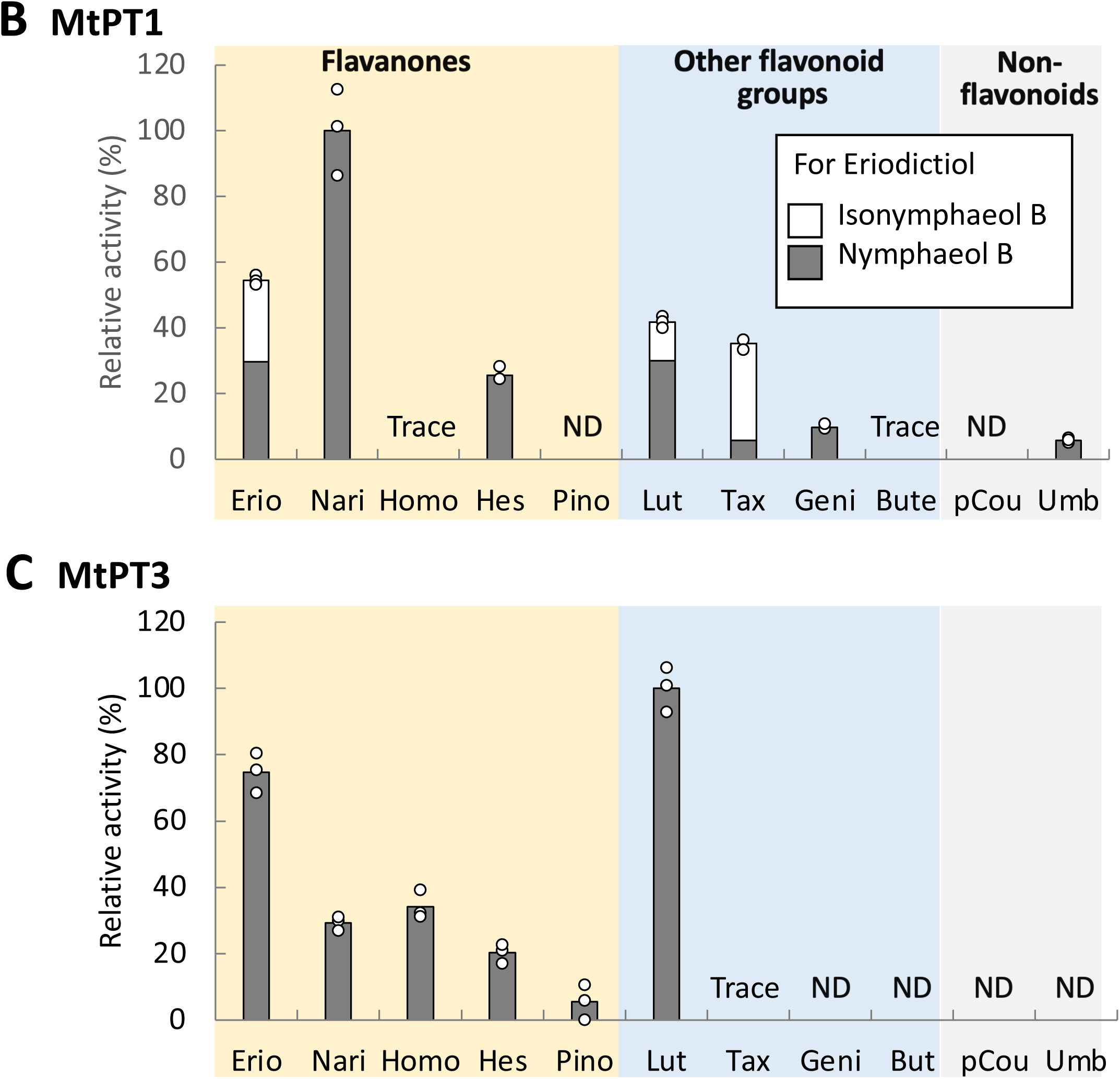
Prenyl acceptor specificity of MtPT1 and MtPT3. (A) The chemical structures of the aromatic molecules tested as prenyl acceptor substrates. Geranyltransferase activities of MtPT1 (B) and MtPT3 (C). The products were quantified as equivalents to the corresponding substrates. The activities less than 5 % of the highest activity were considered trace levels. Multiple activities were combined to form a bar for each substrate, and the values are shown relative to the average of the highest activities. Different colors indicate different products. Independent triplicate reactions were performed for all the aromatic substrates. ND, not detected.

In contrast to naringenin, the removal of all hydroxyl moieties from the B-ring of eriodictyol completely abolished the enzyme activity (pinocembrin) (Figure 7B). Decreases in geranylation activity were also observed for other flavanone molecules with substitution patterns different from that of eriodictyol (homoeriodictyol and hesperitin). This enzyme also accepted molecules belonging to other flavonoid groups at comparable or lower levels than eriodictyol. Furthermore, non-flavonoid molecules were also tested. As a result, a coumarin derivative, umbelliferone, was geranylated by MtPT1, with a simple phenylpropane molecule, *p*-coumaric acid, being unacceptable.

The same prenyl acceptor set was tested for MtPT3 (Figure 7A and C). Among the flavanone molecules tested, MtPT3 showed the highest activity for eriodictyol. This enzyme also accepted the flavone molecule luteolin at a high conversion efficiency (Supplmentary Figure S6) but almost no activities for the other non-flavanone molecules.

Subsequently, prenyl donor specificity was tested for both enzymes using prenyl diphosphates with different chain lengths up to C20 and eriodictyol as a prenyl acceptor. As a result, MtPT1 accepted GPP as the best prenyl donor with FPP accepted at a trace amount equivalent to the control (Figure 8A), while, unexpectedly, MtPT3 preferred FPP to GPP (Figure 8B and Supplementary Figure S7). Considering that MtPT3 is suggested to function in plastids, FPP is not the physiological substrate for this enzyme in *M. tanarius* cells.

**Figure 8.**
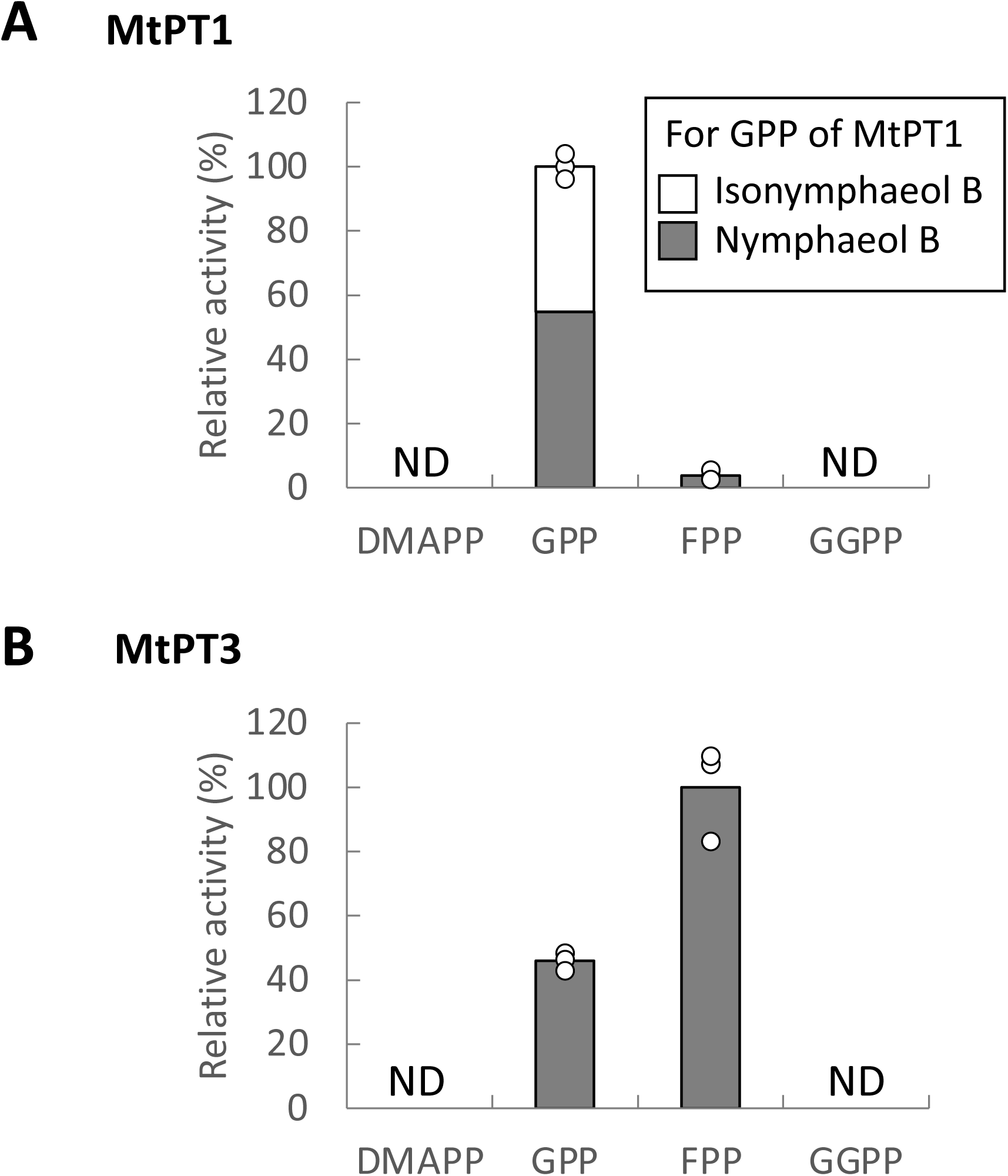
Prenyl donor specificity of MtPT1 and MtPT3. Eriodictyol PT activities of MtPT1 (A) and MtPT3 (B) usinig DMAPP, GPP, FPP, and GGPP as prenyl donor substrates. Independent triplicate reactions were performed for all the prenyl donor substrates. The activities less than 5 % of the highest activity were considered trace levels. Multiple activities were combined to form a bar for each substrate, and the values are shown relative to the average of the highest activities. ND, not detected.

### Synthesis of nymphaeol C by ordered sequential di-prenylation of MtPT1 and MtPT3

*M. tanarius* gulandular trichomes accumulate nymphaeol C, a di-prenylated eriodictyol that possesses one geranyl moiety at the 2′-position of the B-ring and one dimethylallyl moiety at the 6-position of the A-ring. Our biochemical analysis suggested that MtPT1 is responsible for 2′-geranylation, whereas neither MtPT1 nor MtPT3 catalyzed 6-dimethylallylation of the flavanone molecule. To elucidate the biosynthetic pathway of nymphaeol C, we first attempted an all-in-one reaction that contained both MtPT1 and MtPT3 as crude enzymes, eriodictyol as a prenyl acceptor substrate, both DMAPP and GPP as prenyl donor substrates, and Mg^2+^ as a cofactor. LC-MS analysis of the incubation mixture demonstrated the synthesis of nymphaeol C together with mono-geranylated eriodictyols (Figure 9A and Supplementary Figure S8), strongly suggesting mono-geranylated eriodictyol served as a prenyl acceptor substrate. Considering the chemical structures of nymphaeols, we evaluated the nymphaeol B: dimethylallyltransferase activity of MtPT1 and MtPT3 individually, indicating that MtPT3 catalyzed 6-dimethylallylation of nymphaeol B to yield nymphaeol C (Figure 9B and Supplementary Figure S8). These experiments strongly suggest that this di-prenylated molecule is produced from eriodictyol through ordered sequential di-prenylation by MtPT1 and MtPT3. Unfortunately, we were unable to quantify the efficiency of the 6-dimethylallylation of MtPT3 because nymphaeol B standard degraded during storage.

**Figure 9.**
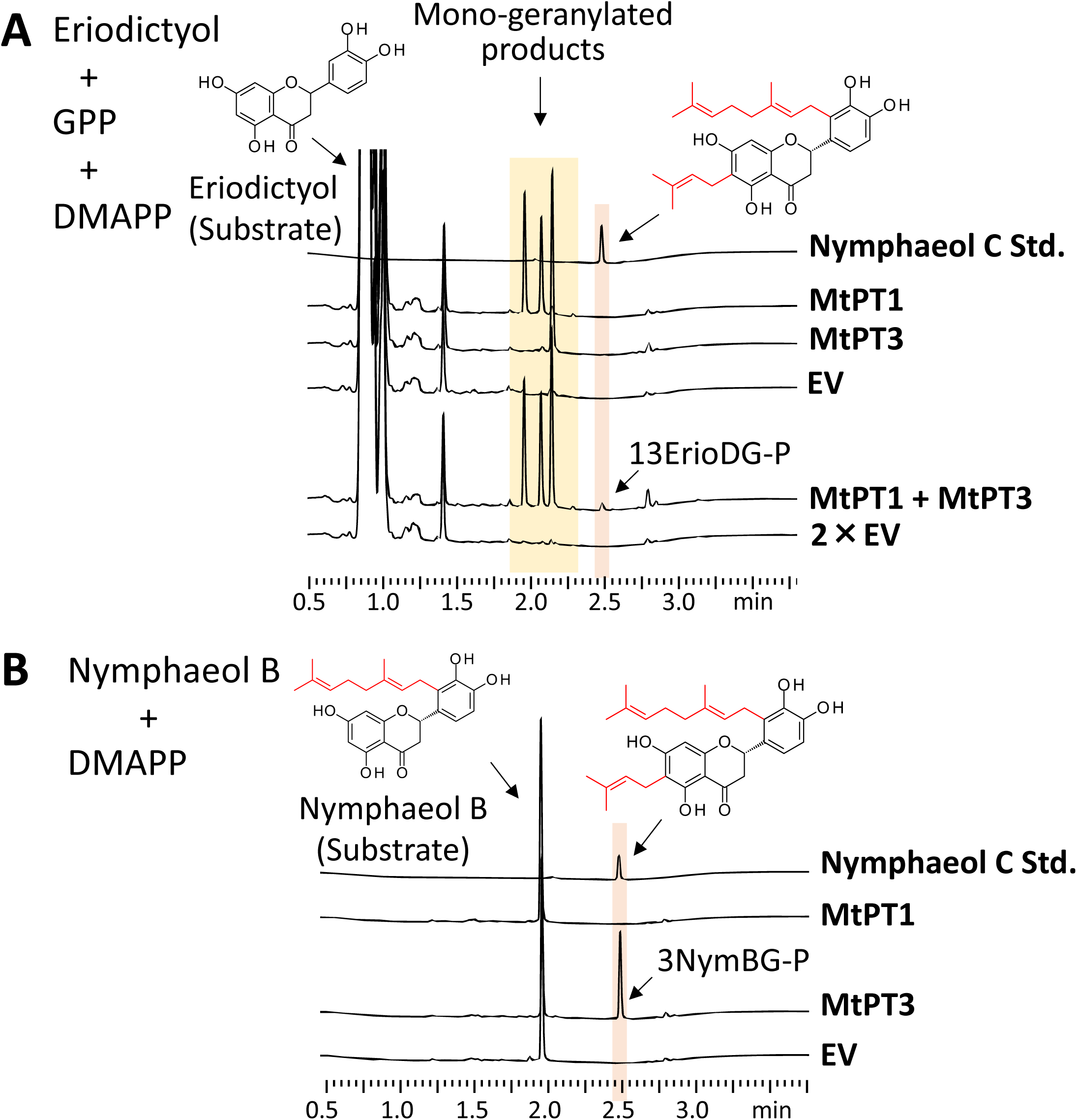
Nymphaeol C synthesis by MtPT1 and MtPT3. (A) Ultraviolet chromatograms (290 nm) of the reaction mixtures in use of eriodictyol as a prenyl acceptor substrate, and both DMAPP and GPP as prenyl donor substrates. Enzyme combinations: MtPT1, MtPT3, and co-existence of MtPT1 and MtPT3. Empty vector was used as a control. (B) Ultraviolet chromatograms (290 nm) of the reaction mixtures of MtPT1 and MtPT3 in use of nymphaeol B and DMAPP as prenyl acceptor and donor substrates, respectively. Empty vector was used as a control.

## Discussion

Nymphaeols have drawn a large attention in the fields of food science and natural medicine because Okinawan propolis exhibited strong antioxidant activity (Kumazawa et al., 2007). In fact, propolis is a natural material that honeybees collect from a waxy part of a plant, e.g. leaf buds *Baccharis dracunculifolia* (Asteraceae), to seal gaps in their hives (Toreti et al., 2013). Because plant species differ depending on the location where beehives are settled, the contents of the propolis largely divergent. Among the various types of domestic propolis in Japan, Okinawan propolis showed one of the strongest antioxidant activities (Kumazawa et al., 2007), and the active components were identified as nymphaeols (Kumazawa et al., 2014). The unique feature of this flavonoid group is the existence of geranyl moiety at the B-ring of eryodictiol. In nature, the most common prenyl moiety attached to the flavonoid core is dimethylallyl, while geranyl is fairly rare. In this study, we were able to characterize a unique PT specific to geranyl, and we also characterized the first secondary metabolite PT from the Euphorbiaceae family (Supplementary Figure S9).

In plant ecology field, *M. tanarius* is well known as a myrmecophilic plant. Myrmecophilic plants live in symbiosis with specific ant species by providing a part of their body as a nesting site for those ants. The *Macaranga* genus includes many species of myrmecophilic plants that form such absolute symbiotic relationships (Fiala et al., 1994). Myrmecophilic plants like *M. tanarius* do not only provide nesting sites for their symbiotic ants but also supply them with food by “food bodies” that develop on the leaf surface. In turn, the symbiotic ants use these food bodies as their primary food source, defend the plants from herbivores, and live exclusively on the plants (Federle et al., 2002). The majority of ant species that form symbiotic relationships with *Macaranga* species belong to the genus *Crematogaster* (Myrmicinae) (Inui et al., 2001). In the *M. tanarius*-*Crematogaster* system, it has been found that a single ant plant species forms a symbiotic relationship with only one or two *Crematogaster* species—which are specialized to that species alone or to a small group of species including its close relatives—thereby creating and maintaining a high degree of species specificity. However, at present, nothing is known about the role that the nymphaeols produced by *M. tanarius* play in their symbiotic relationship with ants.

Because *M. tanarius* is dioecious, it requires pollinators to produce fruit where nymphaeols are specifically accumulated. However, even when the flowers are in full bloom, no honeybees visiting the flowers are seen. It was reported that the most abundant flower visitors found on the male and female inflorescences were *Orius atratus* (Anthocoridae, Hemiptera), a small flower bug (Ishida et al., 2009). This plant has a unique characteristic of direct interactions with three different insects, honeybees, ants, and flower bugs, by which three different biological interactions are working.

Glandular trichomes are often characterized by the accumulation of bioactive secondary metabolites. In herbs of Laminaceae, such as peppermint, basil, and thyme, many monoterpene compounds are exclusively accumulated in glandular trichomes developed on the surface of leaves (Lange, 2015). In hops (Cannabiaceae), bitter acids, which are prenylated phologoglucinol derivatives, and xanthohumol, a prenylated chalcone also known to be accumulated in glandular trichomes of female flowers called lupulins (Tsurumaru et al., 2012). An anti-malaria drug artemisinin, a sesquiterpene lactone, is also accumulated in a limited number of glandular trichomes in *Artemisia annua* (Compositae) (Chen et al., 2017). There appears to be a specific reason why such monoterpenes and prenylated polyphenols are accumulated specifically in glandular trichomes. Because those terpene/polyphenol compounds often exhibit strong biological activities, they are toxic to the producer plants themselves. Then, these active compounds are then sequestrated in a specific organ, such as the glandular trichome, to protect other cells and tissues, such as mesophyll cells. Glandular trichomes therefore keep the toxin safely contained under normal conditions, whereas once the leaves or flowers are attacked by herbivores glandular trichomes are destroyed to release active compounds to repel the enemies.

Aromatic PTs play important roles in the chemical diversification of plant secondary metabolites. After the first discovery of secondary metabolic PT for *p*-hydroxybenzoic acid from *Lithospermum erythrorhizon* (Yazaki et al., 2002), followed by a flavonoid-specific PT identified in *Sophora flavescens* (Sasaki et al., 2008), research on this enzyme family has made significant progress in the last two decades (Munakata and Yazaki, 2024). Thus far, many divergent PTs that accept different aromatic prenyl acceptor substrates have been identified in a variety of plant species. Recently, the discovery of *O*-PTs has also been reported in addition to the conventional *C*-PTs (Munakata et al., 2021; Matsushita et al., 2026). In most cases, those plant PTs show strict specificity for both the substrate molecule and prenylation position. The PTs from *M. tanarius* demonstrated in this study, however, exhibit broad specificity for substrates as a rare example. In terms of prenylation position specificity, MtPT1 and MtPT3 showed distinct preferences for the flavonoid ring each other. Recently, owing to the development of the AlphaFold system, catalytic mechanisms behind the regio-specificity of the membrane-bound PT family have been proposed by combination of mutagenesis and three-dimensional modeling analysis (Huang et al., 2024; Han et al., 2025; Liu et al., 2025). Since screening for the key amino acid(s) in these reports largely relies on comparing PT pairs with different regio-specificities each other, the MtPT1/3 pair that shows a relatively high amino acid identity of 64 % would be a beneficial model to elucidate the recognition mechanism of specific flavonoid rings. Another notable finding is that MtPT3 transfers a dimethylallyl moiety to nymphaeol B (a mono-geranylated eriodictyol) but not to eriodictyol, demonstrating an example that a PT changes prenyl donor acceptance depending on the prenyl acceptor substrate. To date, the prenyl donor substrate specificity of most UbiA PTs has only been assessed using single prenyl acceptor molecules. More comprehensive substrate specificity data for each aromatic PT will enable to evaluate how much the trait of MtPT3 is unique among the UbiA family.

As MtPT1 and MtPT3 form an independent clade in the phylogenetic tree, it is possible that these enzymes have occurred in a manner specific to the taxonomical group, *e.g.*, *Macaranga* or Euphorbiaceae. Recently, genome sequences of Euphorbiaceae species including *M. tanarius* have been published (Liu et al., 2019). These datasets will promote a detailed molecular evolutionary pathway for the acquisition of the aforementioned unique enzymatic functions of MtPT1 and MtPT3.

In *M. tanarius*, these enzymes are suggested to be responsible for the biosynthesis of nymphaeol C, a di-prenylated aromatic molecule. In plants, prenylation modes that form multi-prenylated aromatics differ among plant lineages. In Asteraceae, artepillin C, a bioactive phenylpropane derivative, is formed by di-prenylation of *p-*coumaric acid by a single PT (Munakata et al., 2019). In biosynthesis of hyperforin, a phloroglucinol molecule with four dimethylallyl moieties, in Hypericaceae, each prenylation step is catalyzed by a different PT (Wu et al., 2024). The prenylation mode of nymphaeol C is similar to that of hyperforin, as multiple enzymes are involved in each prenylation step.

In this study, we examined the histological structure of glandular trichomes of *M. tanarius* fruit, in which high amounts of geranylated flavonoids (nymphaeols) are accumulated. From the glandular trichome, we isolated a geranyltransferase (MtPT1) for flavonoids, which is a B-ring-specific PT for eriodyctiol, and showed its broad product specificity. The other PT (MtPT3) is a geranyltransferase for the A-ring with more strict product specificity. MtPT3 accepts DMAPP as a prenyl donor substrate for the B-ring geranylated flavonoid nymphaeol B, while this is not its donor substrate for its non-prenylated form. These are the first PTs in Euphorbiaceae.

## Data availability

The nucleotide sequences of *MtPT1–3* are available in NCBI under the accession numbers LC948606, LC948609 and LC948607, respectively.

## Acknowledgments

We thank Mr. Tsuyoshi Miyagi, Okinawa Prefectural Forest Resources Research Center, Japan for plant identification and the local harvest of raw materials. We thank Dr. Takashi Aoyama of Kyoto University for technical assistance of VP-SEM. We are grateful to Dr. Tomohisa Kuzuyama (The University of Tokyo) and Dr. Takashi Kawasaki (Niigata University) for GPP, Dr. Hirobumi Yamamoto (Toyo University) for DMAPP. These chemicals were used in preliminary experiments. We also thank Dr. David Baulcombe for the pBIN61-P19 plasmid; Dr. Tsuyoshi Nakagawa (Shimane University) for the pGWB505 vector; Dr. Hiroshi Kouchi (International Christian University) for the pHKN29 plasmid; Mr. Yoshiaki Date, and Mr. Shuhei Matsushita, Ms. Kaori Kanazawa (Kyoto University), Dr. Ichino Takuji (Kobe Pharmaceutical University) for technical assistance. Plants were grown in collaboration with the Development and Assessment of Sustainable Humanosphere of the Research Institute for Sustainable Humanosphere (Kyoto University).

## Author contribution

S.K., S.F., K.Y. conceived research. R.M. A.S. K.Y. supervised research, Y.M., R.S., R.M. performed experiments. S.K., S.F. provided authentic compounds and plant materials, respectively. R.M. and K.Y. wrote the manuscript with contribution of all the authors.

## Abbreviations

PT: prenyltransferase
DMAPP: dimethylallyl diphosphate
GPP: geranyl pyrophosphate
MEP: methylerythritol phosphate
PCR: polymerase chain reaction
GFP: green fluorescent protein
LC-PDA: liquid chromatography-photodiode array detector
UPLC: ultra performance liquid chromatography
LC/MS^2^: liquid chromatography / tandem mass spectrometry

## Reference

Ameer B, Weintraub RA, Johnson JV, Yost RA, Rouseff RL (1996) Flavanone absorption after naringin, hesperidin, and citrus administration. Clin Pharmacol Ther 60: 34–40

Bradford MM (1976) A rapid and sensitive method for the quantitation of microgram quantities of protein utilizing the principle of protein-dye binding. Anal Biochem 72: 248–254

Chen M, Yan T, Shen Q, Lu X, Pan Q, Huang Y, Tang Y, Fu X, Liu M, Jiang W, et al (2017) Glandular trichome-specific WRKY1 promotes artemisinin biosynthesis in *Artemisia annua*. New Phytol 214: 304–316

Federle W, Maschwitz U, Hölldobler B (2002) Pruning of host plant neighbours as defence against enemy ant invasions: *Crematogaster* ant partners of *Macaranga* protected by “wax barriers” prune less than their congeners. Oecologia 132: 264–270

Fiala B, Grunsky H, Maschwitz U, Linsenmair KE (1994) Diversity of ant-plant interactions: protective efficacy in *Macaranga* species with different degrees of ant association. Oecologia 97: 186–192

Guindon S, Dufayard J-F, Lefort V, Anisimova M, Hordijk W, Gascuel O (2010) New algorithms and methods to estimate maximum-likelihood phylogenies: assessing the performance of PhyML 3.0. System Biol 59: 307–321

Hallgren J, Tsirigos KD, Pedersen MD, Almagro Armenteros JJ, Marcatili P, Nielsen H, Krogh A, Winther O (2022) DeepTMHMM predicts alpha and beta transmembrane proteins using deep neural networks. biorxiv 2022–04

Han J, Munakata R, Takahashi H, Koeduka T, Kubota M, Moriyoshi E, Hehn A, Sugiyama A, Yazaki K (2025) Catalytic mechanism underlying the regiospecificity of coumarin-substrate transmembrane prenyltransferases in Apiaceae. Plant Cell Physiol 66: 1–14

Huang X-C, Tang H, Wei X, He Y, Hu S, Wu J-Y, Xu D, Qiao F, Xue J-Y, Zhao Y (2024) The gradual establishment of complex coumarin biosynthetic pathway in Apiaceae. Nat Commun 15: 6864

Inui Y, Itioka T, Murase K, Yamaoka R, Itino T (2001) Chemical recognition of partner plant species by foundress ant queens in *Macaranga*–*Crematogaster* myrmecophytism. J Chem Ecol 27: 2029–2040

Ishida C, Kono M, Sakai S (2009) A new pollination system: brood-site pollination by flower bugs in *Macaranga* (Euphorbiaceae). Annal Bot 103: 39–44

Kalyaanamoorthy S, Minh BQ, Wong TK, Von Haeseler A, Jermiin LS (2017) ModelFinder: fast model selection for accurate phylogenetic estimates. Nat Methods 14: 587–589

Karamat F, Olry A, Munakata R, Koeduka T, Sugiyama A, Paris C, Hehn A, Bourgaud F, Yazaki K (2014) A coumarin - specific prenyltransferase catalyzes the crucial biosynthetic reaction for furanocoumarin formation in parsley. Plant J 77: 627–638

Katoh K, Rozewicki J, Yamada KD (2019) MAFFT online service: multiple sequence alignment, interactive sequence choice and visualization. Brief Bioinform 20: 1160–1166

Kumagai H, Kouchi H (2003) Gene Silencing by Expression of hairpin RNA in *Lotus japonicus* roots and root nodules. MPMI 16: 663–668

Kumazawa S, Murase M, Momose N, Fukumoto S (2014) Analysis of antioxidant prenylflavonoids in different parts of *Macaranga tanarius*, the plant origin of Okinawan propolis. Asian Pac J Trop Med 7: 16–20

Kumazawa S, Nakamura J, Murase M, Miyagawa M, Ahn M-R, Fukumoto S (2008) Plant origin of Okinawan propolis: honeybee behavior observation and phytochemical analysis. Naturwissenschaften 95: 781–786

Kumazawa S, Ueda R, Hamasaka T, Fukumoto S, Fujimoto T, Nakayama T (2007) Antioxidant prenylated flavonoids from propolis collected in Okinawa, Japan. J Agric Food Chem 55: 7722–7725

Kuraku S, Zmasek CM, Nishimura O, Katoh K (2013) aLeaves facilitates on-demand exploration of metazoan gene family trees on MAFFT sequence alignment server with enhanced interactivity. Nucleic Acids Res 41: W22–W28

Lange BM (2015) The evolution of plant secretory structures and emergence of terpenoid chemical diversity. Annu Rev Plant Biol 66: 139–159

Leveson - Gower RB, Roelfes G (2022) Biocatalytic Friedel - Crafts Reactions. ChemCatChem 14: e202200636

Liu H, Wei J, Yang T, Mu W, Song B, Yang T, Fu Y, Wang X, Hu G, Li W (2019) Molecular digitization of a botanical garden: high-depth whole-genome sequencing of 689 vascular plant species from the Ruili Botanical Garden. GigaScience 8: giz007

Liu S, Tao Y, Zhang Y, Gong J, Wu Z, Wang Z, An Z, Shi R, Zhao Y, Shawky E, et al (2025) Identification, Characterization, and catalytic mechanism of regioselective UbiA prenyltransferases in *Morus* plants. Angew Chem 137: e202504190

Matsushita S, Munakata R, Roumani M, Olry A, Nakayasu M, Hehn A, Matsukawa T, Sugiyama A, Yazaki K (2026) A membrane-bound aromatic *O*-prenyltransferase catalyzes the last reaction step in citrus auraptene biosynthesis. *bioRxiv* 2026–07

Minh BQ, Nguyen MAT, Von Haeseler A (2013) Ultrafast approximation for phylogenetic bootstrap. Mol Biol Evol 30: 1188–1195

Munakata R, Inoue T, Koeduka T, Karamat F, Olry A, Sugiyama A, Takanashi K, Dugrand A, Froelicher Y, Tanaka R (2014) Molecular cloning and characterization of a geranyl diphosphate-specific aromatic prenyltransferase from lemon. Plant Physiol 166: 80–90

Munakata R, Kitajima S, Nuttens A, Tatsumi K, Takemura T, Ichino T, Galati G, Vautrin S, Bergès H, Grosjean J, et al (2020) Convergent evolution of the UbiA prenyltransferase family underlies the independent acquisition of furanocoumarins in plants. New Phytol 225: 2166–2182

Munakata R, Olry A, Takemura T, Tatsumi K, Ichino T, Villard C, Kageyama J, Kurata T, Nakayasu M, Jacob F, et al (2021) Parallel evolution of UbiA superfamily proteins into aromatic *O*-prenyltransferases in plants. Proc Natl Acad Sci USA 118: e2022294118

Munakata R, Takemura T, Tatsumi K, Moriyoshi E, Yanagihara K, Sugiyama A, Suzuki H, Seki H, Muranaka T, Kawano N (2019) Isolation of *Artemisia capillaris* membrane-bound di-prenyltransferase for phenylpropanoids and redesign of artepillin C in yeast. Commun Biol 2: 384

Munakata R, Yazaki K (2024) How did plants evolve the prenylation of specialized phenolic metabolites by means of UbiA prenyltransferases? Curr Opin Plant Biol 81: 102601

Nakagawa T, Suzuki T, Murata S, Nakamura S, Hino T, Maeo K, Tabata R, Kawai T, Tanaka K, Niwa Y (2007) Improved Gateway binary vectors: high-performance vectors for creation of fusion constructs in transgenic analysis of plants. Biosci Biotechnol Biochem 71: 2095–2100

Nguyen L-T, Schmidt HA, Von Haeseler A, Minh BQ (2015) IQ-TREE: a fast and effective stochastic algorithm for estimating maximum-likelihood phylogenies. Mol Biol Evol 32: 268–274

Norkunas K, Harding R, Dale J, Dugdale B (2018) Improving agroinfiltration-based transient gene expression in *Nicotiana benthamiana*. Plant Methods 14: 71

Norris SR, Barrette TR, DellaPenna D (1995) Genetic dissection of carotenoid synthesis in Arabidopsis defines plastoquinone as an essential component of phytoene desaturation. Plant Cell 7: 2139–2149

Ødum MT, Teufel F, Thumuluri V, Almagro Armenteros JJ, Johansen AR, Winther O, Nielsen H (2024) DeepLoc 2.1: multi-label membrane protein type prediction using protein language models. Nucleic Acids Res 52: W215–W220

Ohara K, Yamamoto K, Hamamoto M, Sasaki K, Yazaki K (2006) Functional characterization of *OsPPT1*, which encodes *p*-hydroxybenzoate polyprenyltransferase involved in ubiquinone biosynthesis in *Oryza sativa*. Plant Cell Physiol 47: 581–590

Ramos FP., Iwamoto L, Piva VH, Teixeira SP (2024) Updating the Knowledge on the Secretory Machinery of Hops (*Humulus lupulus* L., Cannabaceae) Plants, 13: 864

Sasaki K, Mito K, Ohara K, Yamamoto H, Yazaki K (2008) Cloning and characterization of naringenin 8-prenyltransferase, a flavonoid-specific prenyltransferase of *Sophora flavescens*. Plant Physiol 146: 1075–1084

Sattler SE, Gilliland LU, Magallanes-Lundback M, Pollard M, DellaPenna D (2004) Vitamin E is essential for seed longevity and for preventing lipid peroxidation during germination. Plant Cell 16: 1419–1432

Simons R, Vincken J, Bakx EJ, Verbruggen MA, Gruppen H (2009) A rapid screening method for prenylated flavonoids with ultra - high - performance liquid chromatography/electrospray ionisation mass spectrometry in licorice root extracts. Rapid Comm Mass Spectrometry 23: 3083–3093

Toreti VC, Sato HH, Pastore GM, Park YK (2013) Recent progress of propolis for its biological and chemical compositions and its botanical origin. Evid Based Complement Alternat Med 2013: 1–13

Tsurumaru Y, Sasaki K, Miyawaki T, Uto Y, Momma T, Umemoto N, Momose M, Yazaki K (2012) HlPT-1, a membrane-bound prenyltransferase responsible for the biosynthesis of bitter acids in hops. Biochemi Biophys Res Commun 417: 393–398

Wu S, Morotti ALM, Yang J, Wang E, Tatsis EC (2024) Single-cell RNA sequencing facilitates the elucidation of the complete biosynthesis of the antidepressant hyperforin in St. John’s wort. Mol Plant 17: 1439–1457

Yang T, Fang L, Sanders S, Jayanthi S, Rajan G, Podicheti R, Thallapuranam SK, Mockaitis K, Medina-Bolivar F (2018) Stilbenoid prenyltransferases define key steps in the diversification of peanut phytoalexins. J Biol Chem 293: 28–46

Yang X, Jiang Y, Yang J, He J, Sun J, Chen F, Zhang M, Yang B (2015) Prenylated flavonoids, promising nutraceuticals with impressive biological activities. Trends Food Sci Technol 44: 93–104

Yazaki K, Kunihisa M, Fujisaki T, Sato F (2002) Geranyl diphosphate: 4-hydroxybenzoate geranyltransferase from *Lithospermum erythrorhizon*: cloning and characterization of a key enzyme in shikonin biosynthesis. J Biol Chem 277: 6240–6246

Yazaki K, Sasaki K, Tsurumaru Y (2009) Prenylation of aromatic compounds, a key diversification of plant secondary metabolites. Phytochemistry 70: 1739–1745

