## Supplementary Tables and Figures for "Biosynthesis of prenylated flavonoids by two membrane-bound prenyltransferases in glandular trichomes of *Macaranga tanarius*"

TLC Silicagel 60 F254 0.5mm (MERCK)
Toluene：ethyl acetate：acetic acid＝70：30：0.3

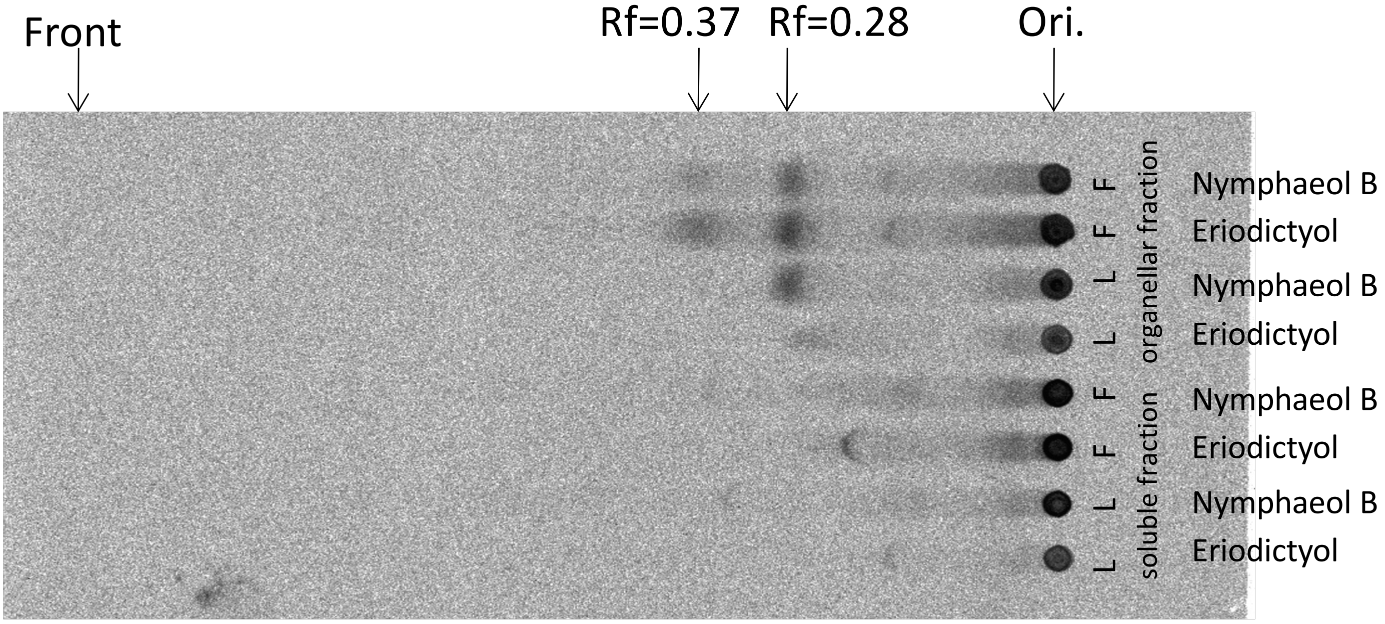

F＝ flower L＝ leaf

**Supplementary Figure 1. Prenyltransferase activity of native crude enzymes from *M. tanarius.***

Cell-free extracts of *M. tanarius* leaves were fractionated into organelle-rich and soluble fractions (supernatant of 100,000 × *g*). In the enzyme assay, $^{14}C$-DMAPP, and eriodictyol and nymphaeol B were used as the prenyl donor and acceptor substrates, respectively. Reaction products were detected by radio-TLC.

Munakata *et al.*

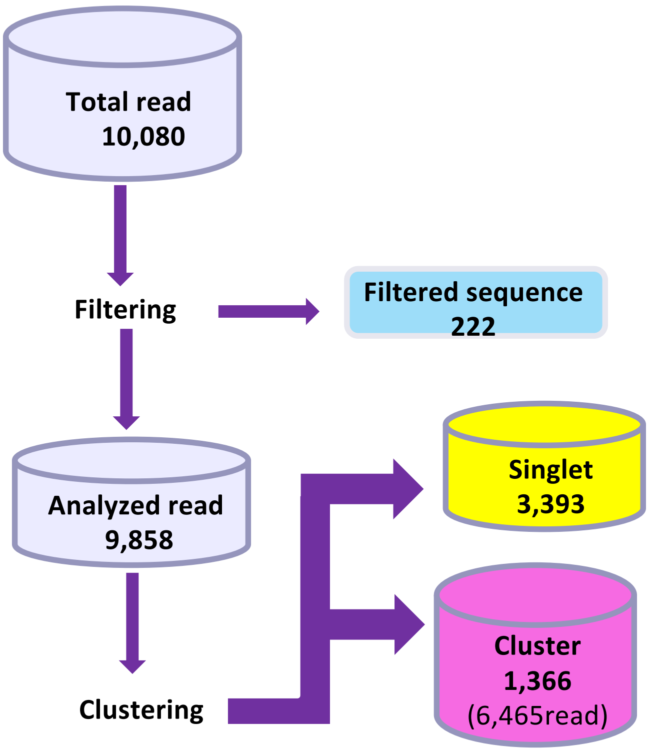

**Supplementary Figure 2. The workflow of EST sequence data analysis**

Munakata *et al*.

**A**

**MtPT1**

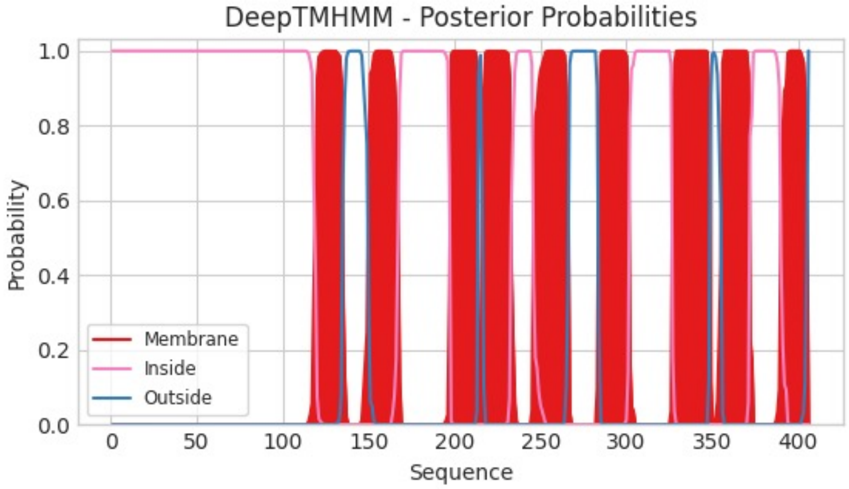

**MtPT2**

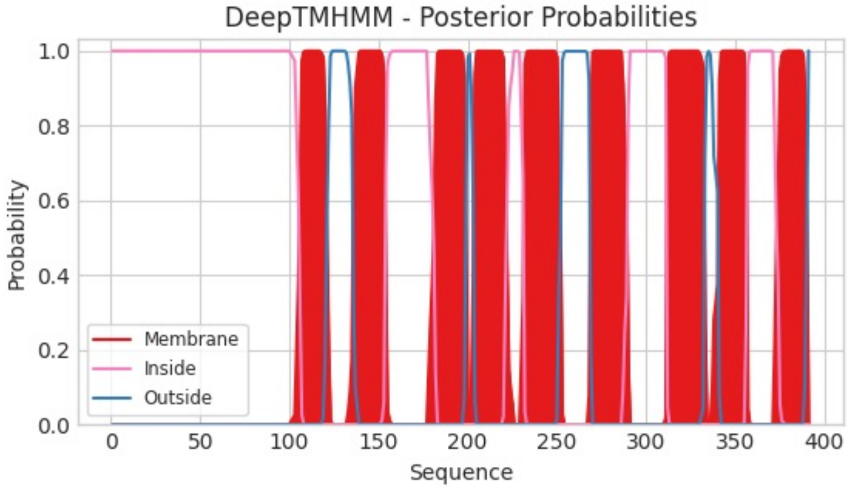

**MtPT3**

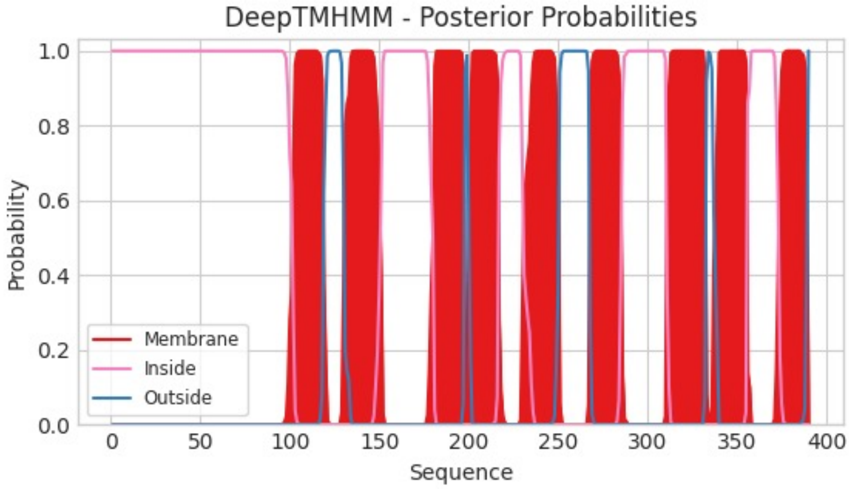

**Supplementary Figure S3. *In silico* analysis of MtPT1–3 polypeptides.**

**B**

| Protein_ID | MtPT1 | MtPT2 | MtPT3 |
|---|---|---|---|
| **Localizations** | Plastid | Plastid | Plastid |
| **Signals** | | | |
| **Cytoplasm** | 0.0867 | 0.095 | 0.0916 |
| **Nucleus** | 0.094 | 0.076 | 0.0849 |
| **Extracellular** | 0.0279 | 0.0141 | 0.0255 |
| **Cell membrane** | 0.1341 | 0.0829 | 0.1259 |
| **Mitochondrion** | 0.1839 | 0.1732 | 0.1163 |
| **Plastid** | **0.9291** | **0.9289** | **0.8918** |
| **Endoplasmic reticulum** | 0.2012 | 0.1363 | 0.1715 |
| **Lysosome/Vacuole** | 0.0645 | 0.1201 | 0.0929 |
| **Golgi apparatus** | 0.1139 | 0.1876 | 0.138 |
| **Peroxisome** | 0.1207 | 0.1085 | 0.1247 |

**Supplementary Figure S3. *In silico* analysis of MtPT1–3 polypeptides.**
***– continued***

**C**

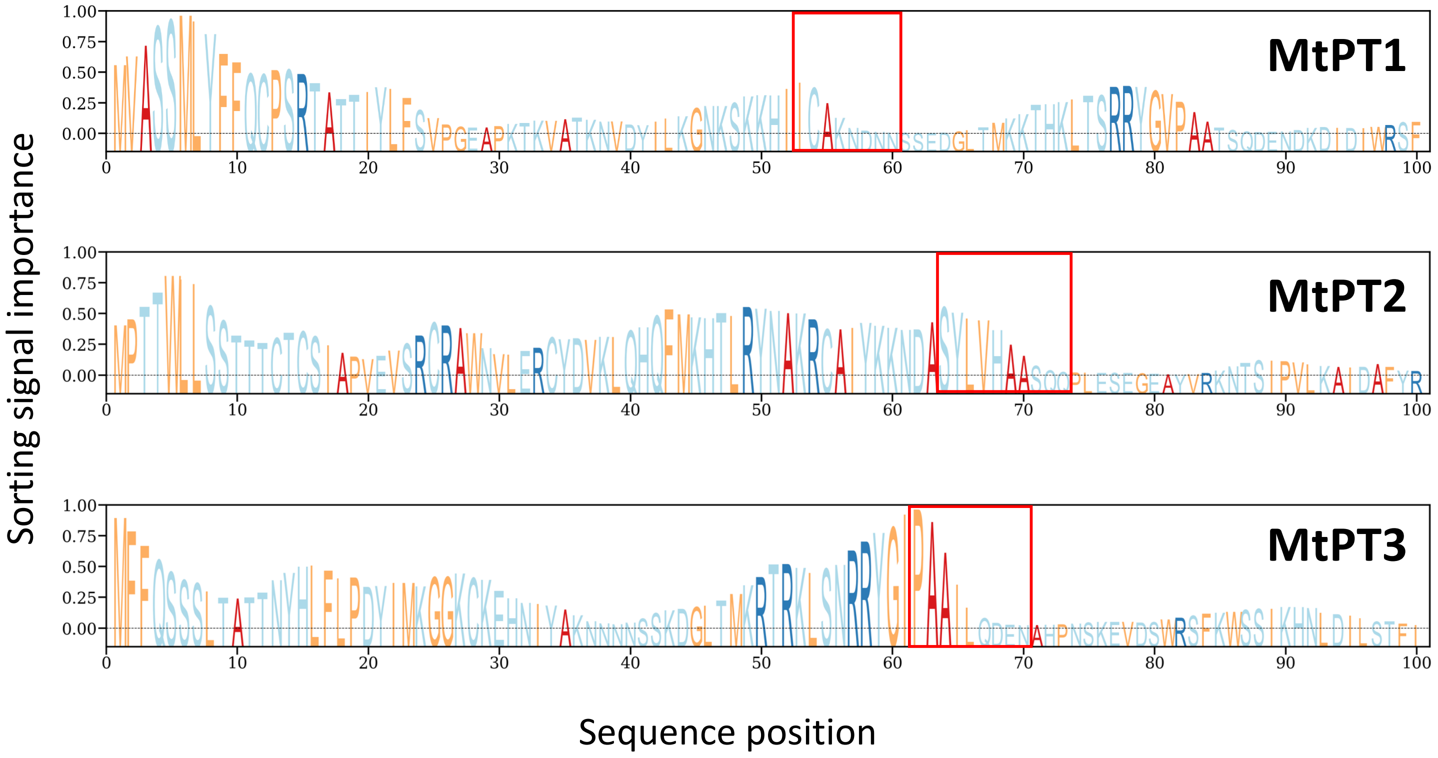

**Supplementary Figure S3.** ***In silico*** **analysis of MtPT1–3 polypeptides. –** ***continued***

(A) Transmembrane regions of MtPT1–3 predicted by DeepTMHMM. The vertical axis represents the possibility of being part of a transmembrane region for each amino acid residue. (B) Subcellular localization of MtPT1–3 predicted by DeepLoc2.1. (C) Prediction of important amino acid regions for a sorting signal predicted by DeepLoc2.1. The red boxes highlight the amino acid regions that possibly indicate the end of the sorting signals.

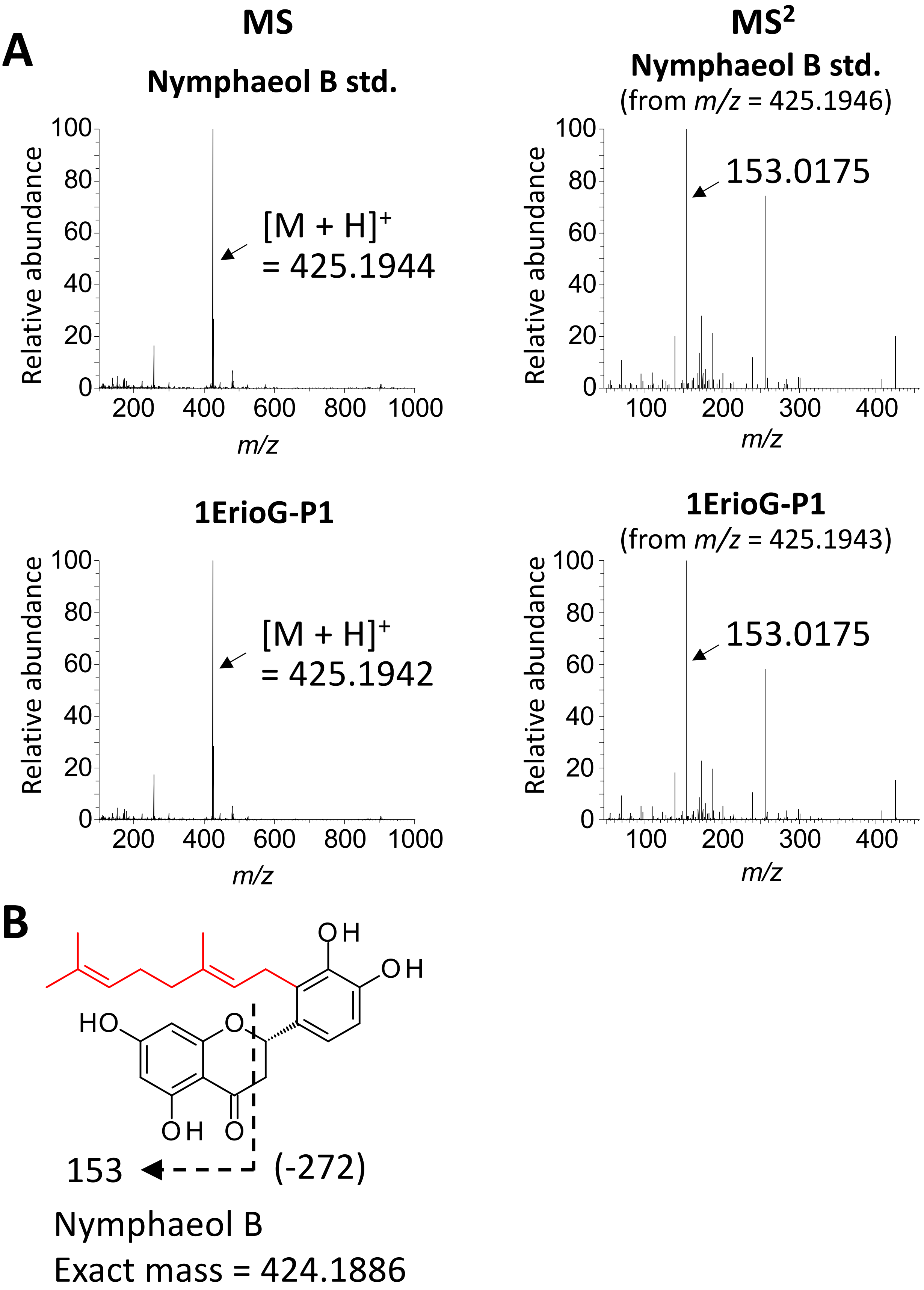

**Supplementary Figure S4. MS$^2$ analysis of enzymatic reaction products of MtPT1 and MtPT3 in eriodictyol geranyltransferase assay**

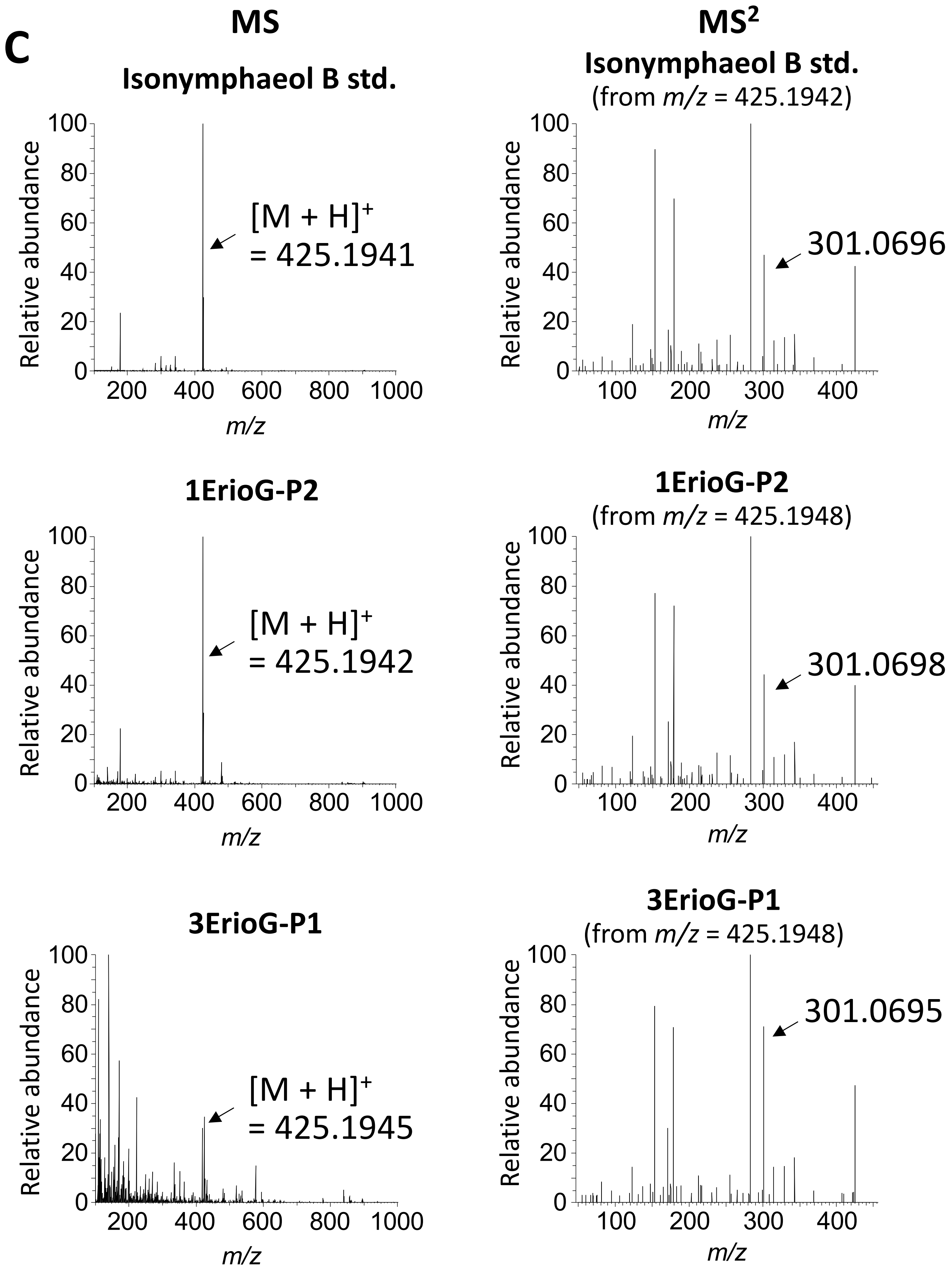

**Supplementary Figure S4. MS² analysis of enzymatic reaction products of MtPT1 and MtPT3 in eriodictyol geranyltransferase assay. – *continued***

Munakata *et al.*

**D**

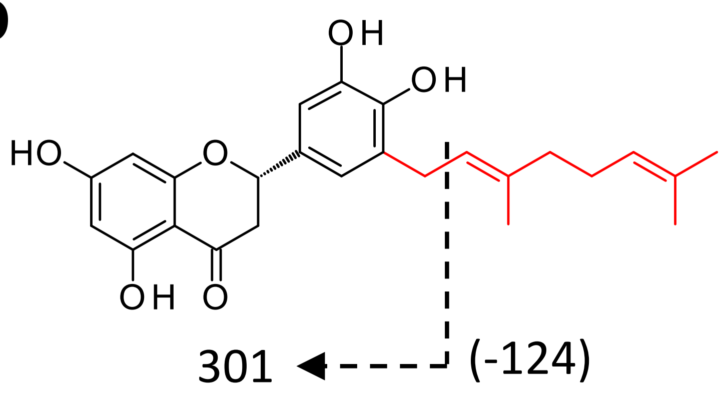

Isonymphaeol B
Exact mass = 424.1886

**Supplementary Figure S4. MS$^2$ analysis of enzymatic reaction products of MtPT1 and MtPT3 in eriodictyol geranyltransferase assay. – *continued***

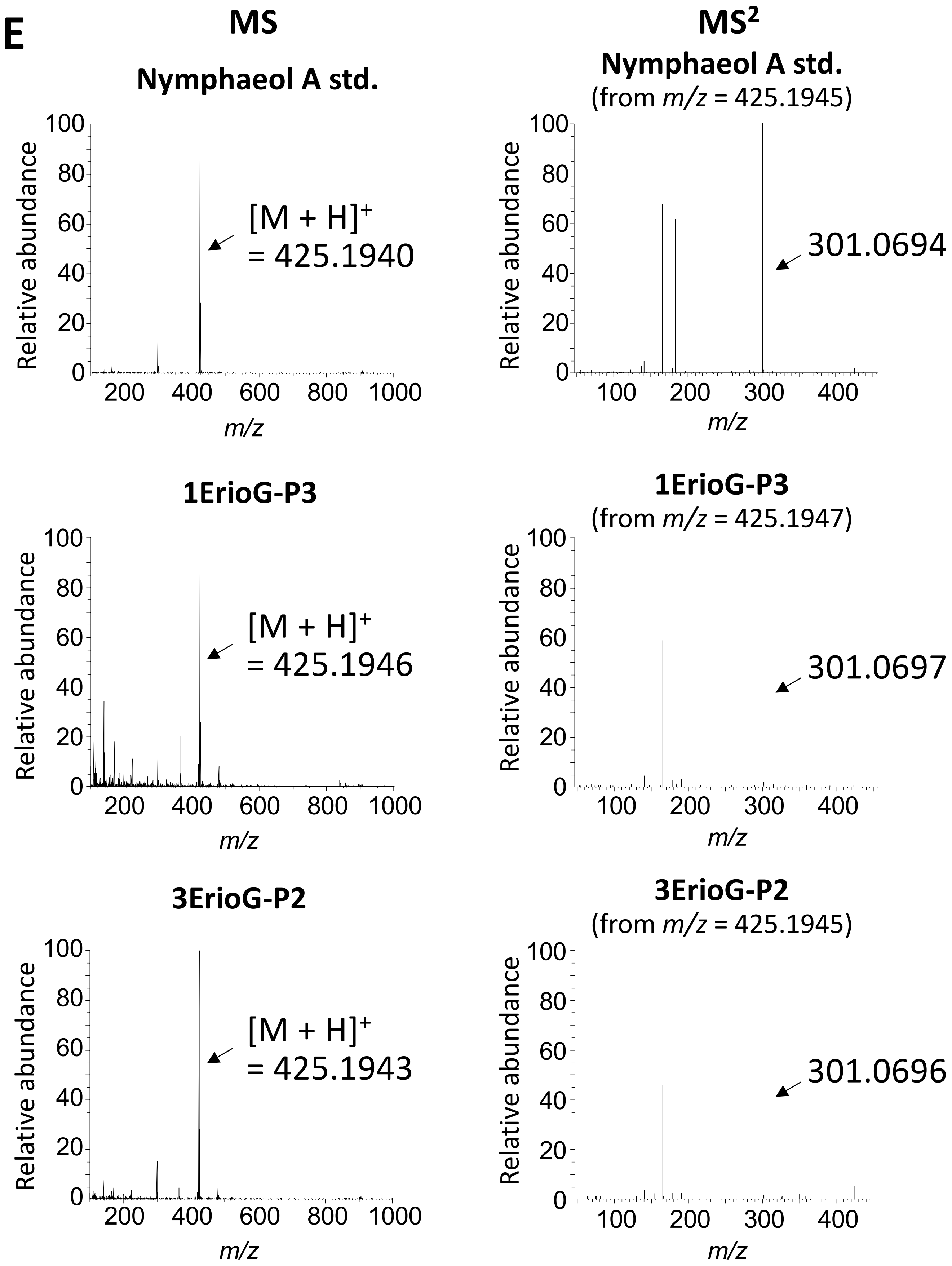

**Supplementary Figure S4. $MS^2$ analysis of enzymatic reaction products of MtPT1 and MtPT3 in eriodictyol geranyltransferase assay. – *continued***

Munakata *et al.*

**F**

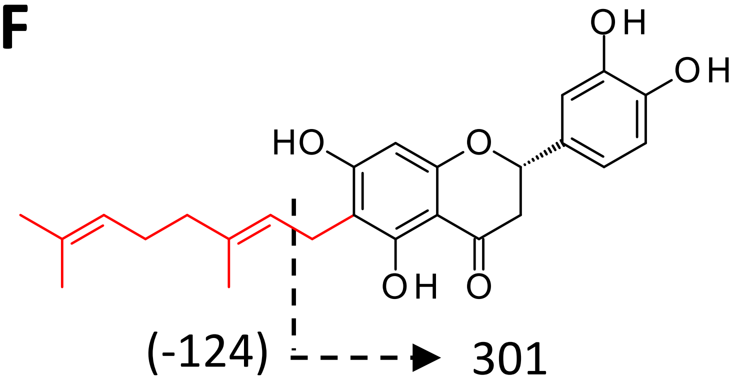

Nymphaeol A
Exact mass = 424.1886

**Supplementary Figure S4. $MS^2$ analysis of enzymatic reaction products of MtPT1 and MtPT3 in eriodictyol geranyltransferase assay. – *continued***

**G**

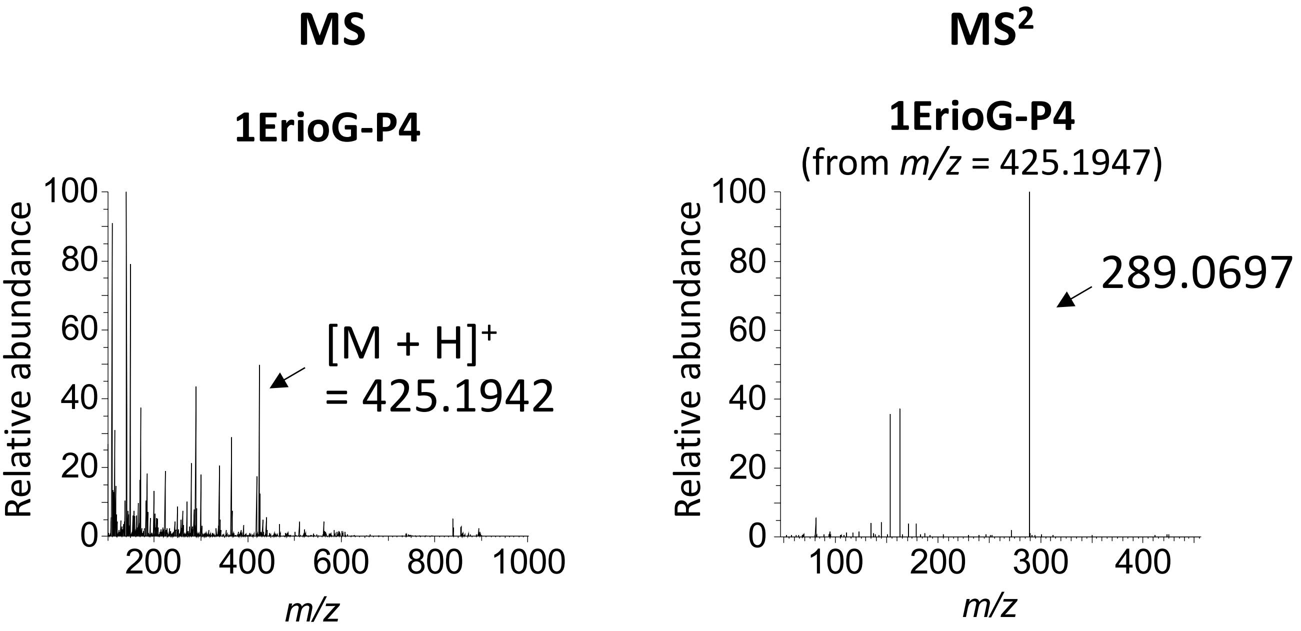

**H**

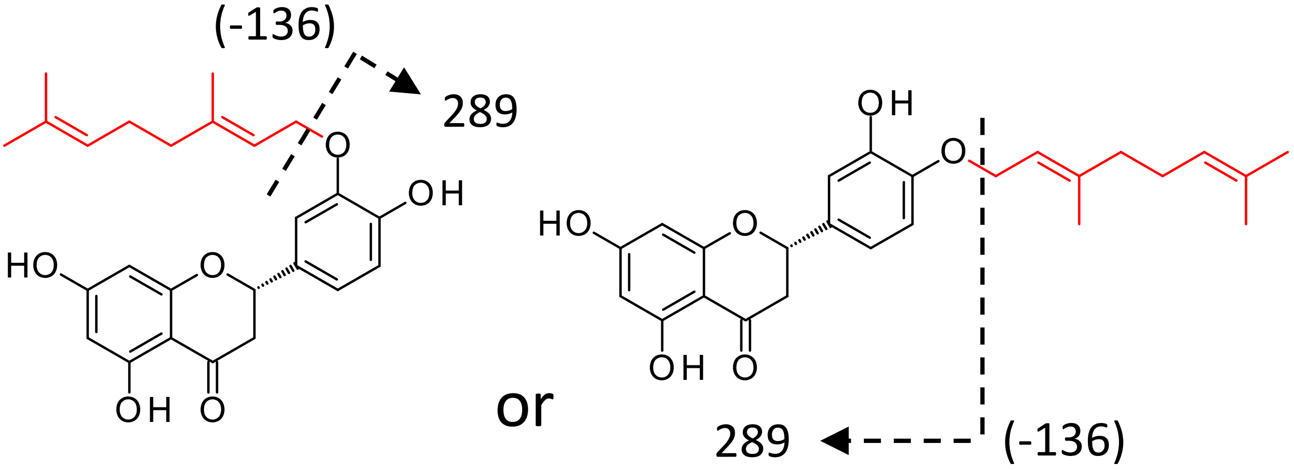

Putative chemical structure
Exact mass = 424.1886

**Supplementary Figure S4. MS² analysis of enzymatic reaction products of MtPT1 and MtPT3 in eriodictyol geranyltransferase assay. – *continued***

Munakata *et al.*

**Supplementary Figure S4. Legend**

(A, C, E, and G) MS (left panels) and $MS^2$ (right panels) spectra of the enzymatic reaction products in the positive ion mode (A, 1ErioG-P1; C, 1ErioG-P2 and 3ErioG-P1; E, 1ErioG-P3 and 3ErioG-P2; G, 1ErioG-P4). The molecular ion peaks and characteristic fragment ions are indicated. (B, D, F, and H) The chemical structure of the reaction products. The exact masses and fragmentation patterns to give the fragment ions indicated in the $MS^2$ spectra are shown. Putative structures are shown in (H) due to the lack of a corresponding standard.

.

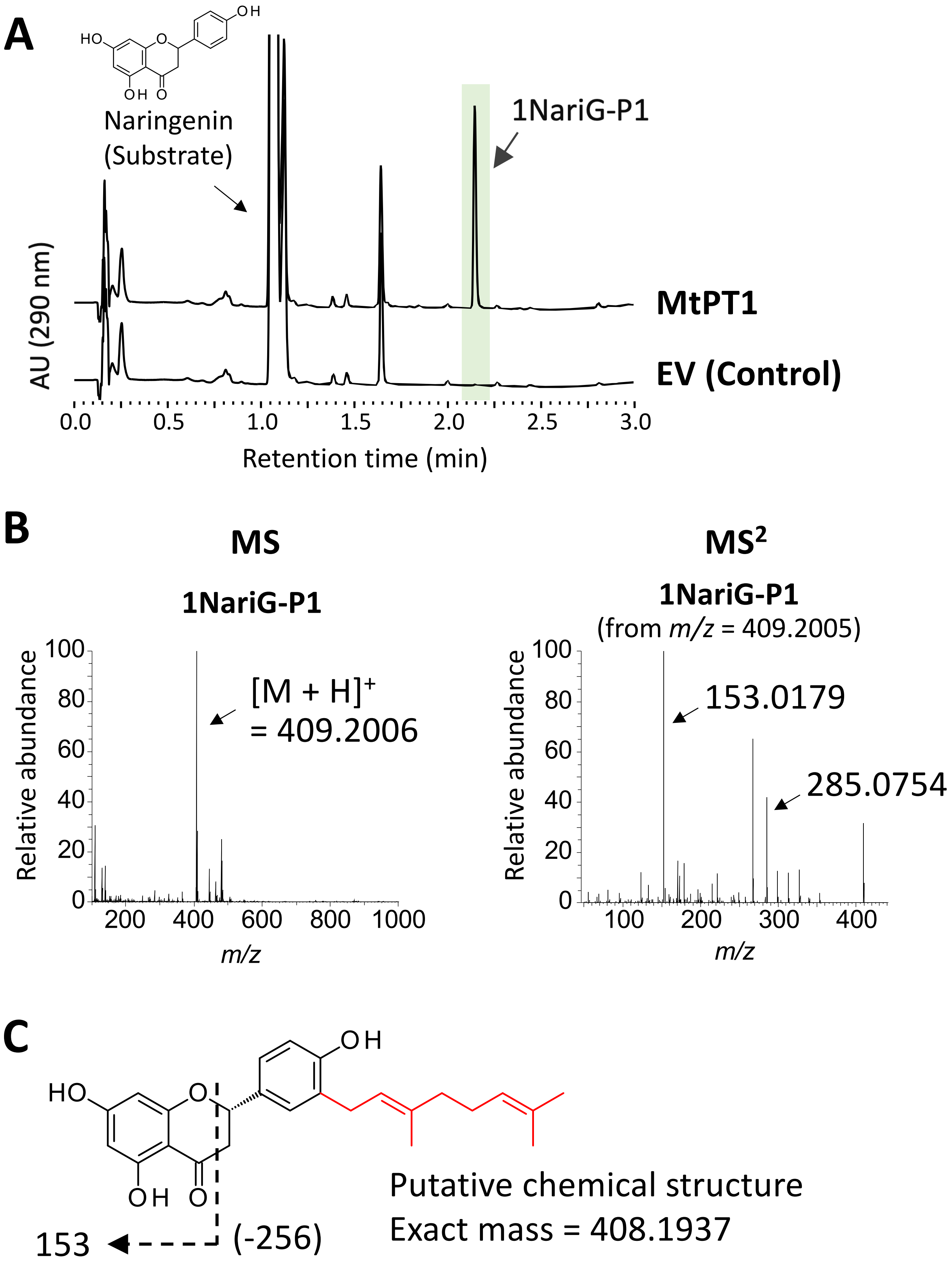

**Supplementary Figure S5. LC-PDA and MS$^2$ analyses of the enzymatic reaction product of MtPT1 in naringenin geranyltransferase assay.**

Munakata *et al.*

**Supplementary Figure S5. Legend**

(A) An ultraviolet chromatogram (290 nm) of the reaction mixture of recombinant MtPT1 in naringenin geranyltransferase assay. Empty vector was used as a control. (B) MS (left panels) and $MS^2$ (right panel) spectra of the enzymatic reaction product 1NariG-P1 in the positive ion mode. The molecular ion peak and characteristic fragment ions are indicated. (C) The putative chemical structure of 1NariG-P1. The exact mass and fragmentation pattern to give the fragment ion indicated in the $MS^2$ spectrum are shown.

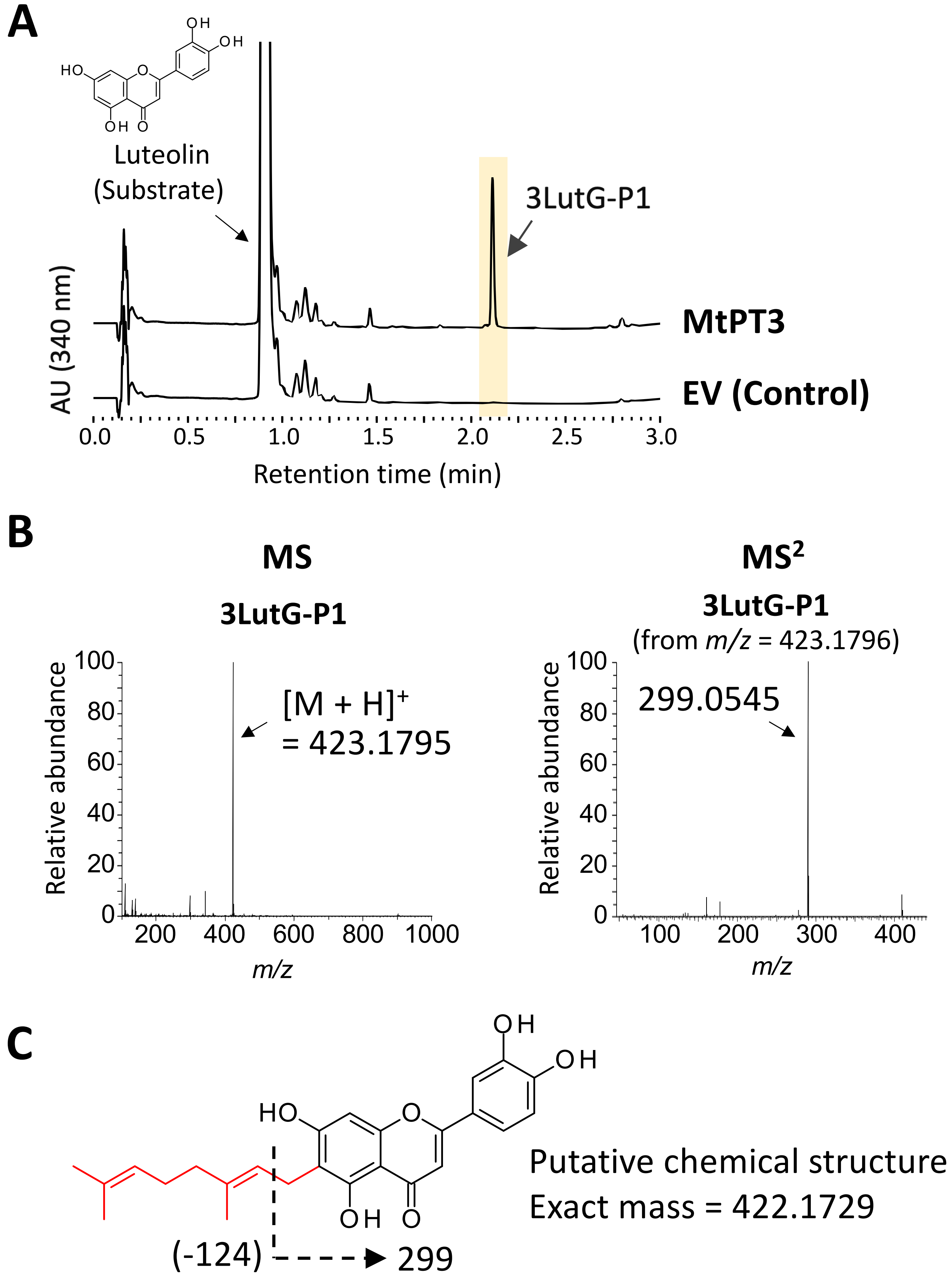

**Supplementary Figure S6. LC-PDA and MS² analyses of the main enzymatic reaction product of MtPT3 in luteolin geranyltransferase assay.**

Munakata *et al.*

**Supplementary Figure S6.** (legend)
(A) An ultraviolet chromatogram (340 nm) of the reaction mixture of recombinant MtPT1 in luteolin geranyltransferase assay. Empty vector was used as a control. (B) MS (left panels) and $MS^2$ (right panel) spectra of the enzymatic reaction product 3LutG-P1 in the positive ion mode. The molecular ion peak and the characteristic fragment ion are indicated. (C) The putative chemical structure of 3LutG-P1. The exact mass and fragmentation pattern to give the fragment ion indicated in the $MS^2$ spectrum are shown.

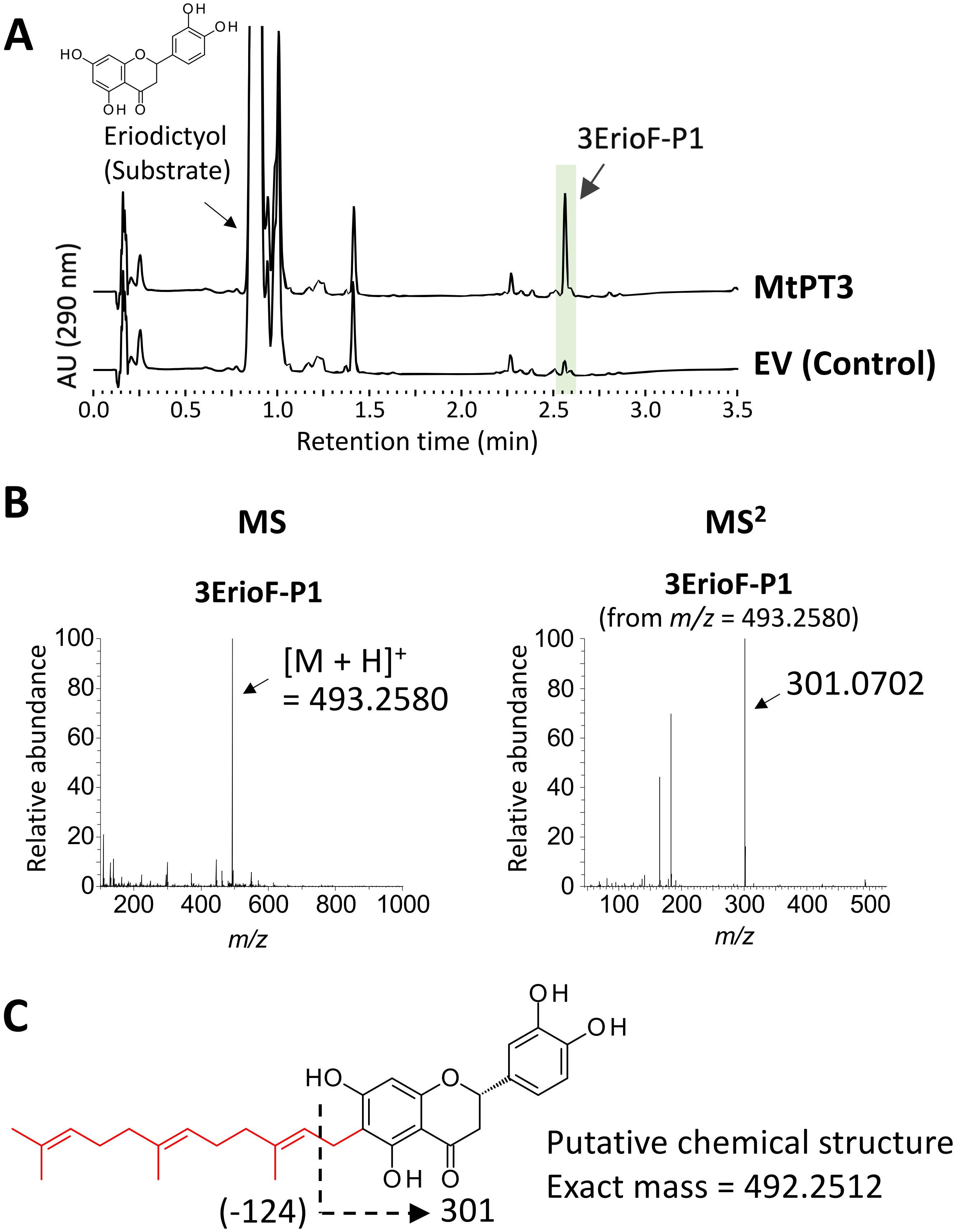

**Supplementary Figure S7. LC-PDA and MS$^2$ analyses of the enzymatic reaction product of MtPT3 in eriodictyol farnesyltransferase assay.**

Munakata *et al.*

***– continued***

(A) An ultraviolet chromatogram (290 nm) of the reaction mixture of recombinant MtPT1 in eriodictyol farnesyltransfarase assay. Empty vector was used as a control. (B) MS (left panels) and $MS^2$ (right panel) spectra of the enzymatic reaction product 3ErioF-P1 in the positive ion mode. The molecular ion peak and the characteristic fragment ion are indicated. (C) The putative chemical structure of 3ErioF-P1. The exact mass and fragmentation pattern to give the fragment ion indicated in the $MS^2$ spectrum are shown.

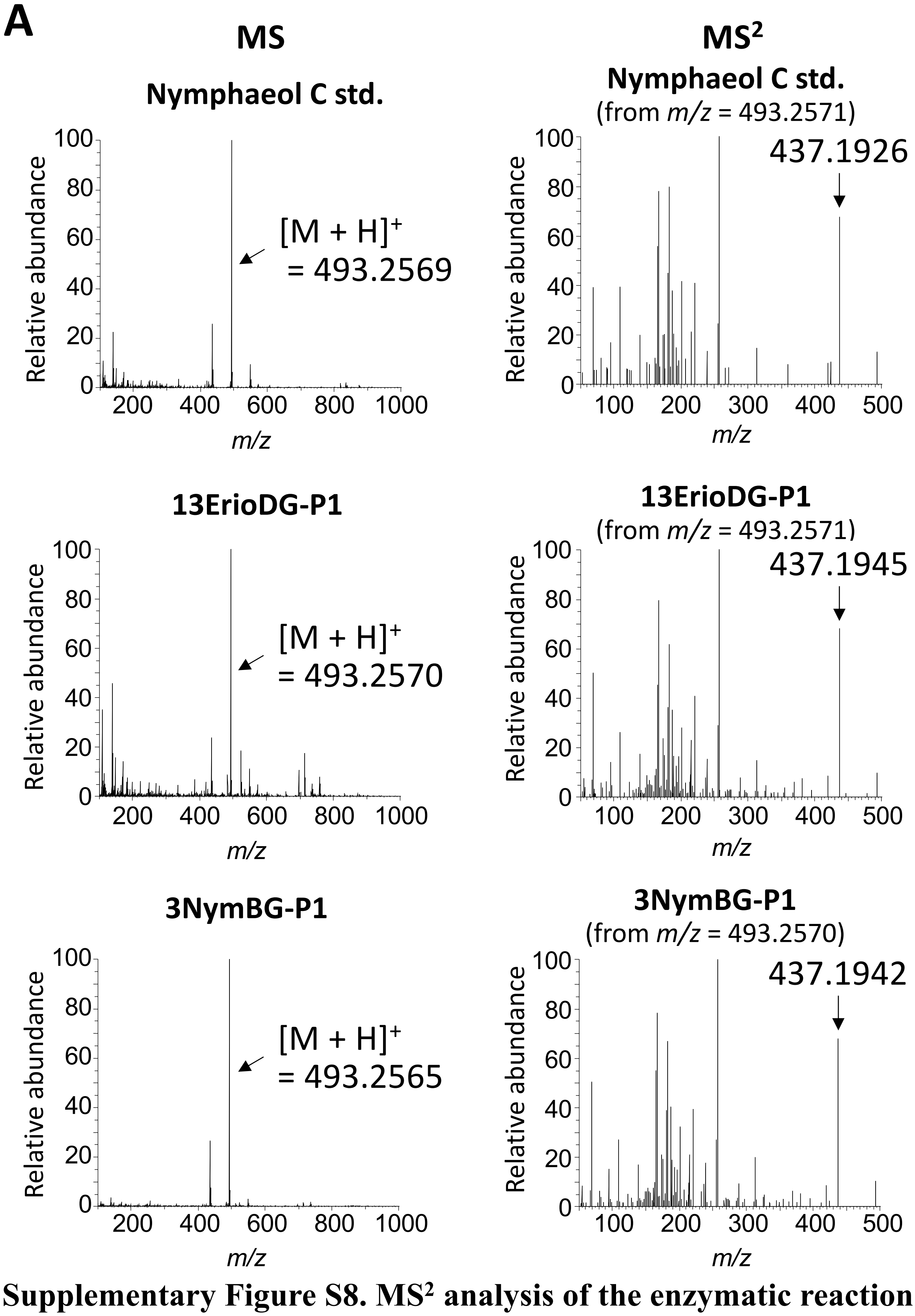

**Supplementary Figure S8. MS$^2$ analysis of the enzymatic reaction product in nymphaeol C synthesis assay.**

**B**

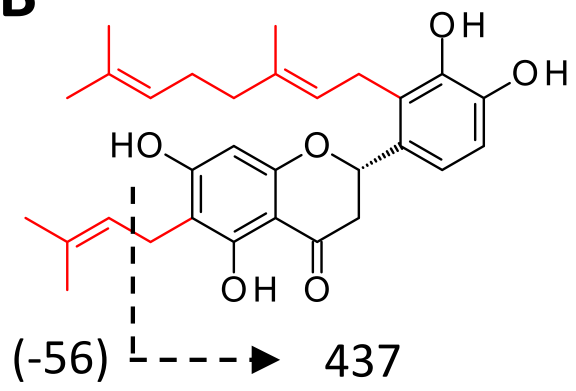

Nymphaeol C
Exact mass = 492.2512

***– continued***

(A) MS (left panels) and MS$^2$ (right panels) spectra of the enzymatic reaction products in the positive ion mode (13ErioDG-P1 and 3NymBG-P1). The molecular ion peaks and the characteristic fragment ions are indicated. (B) The chemical structure of the reaction products. Their exact masses and fragmentation patterns to give the fragment ions indicated in the MS$^2$ spectra are shown.

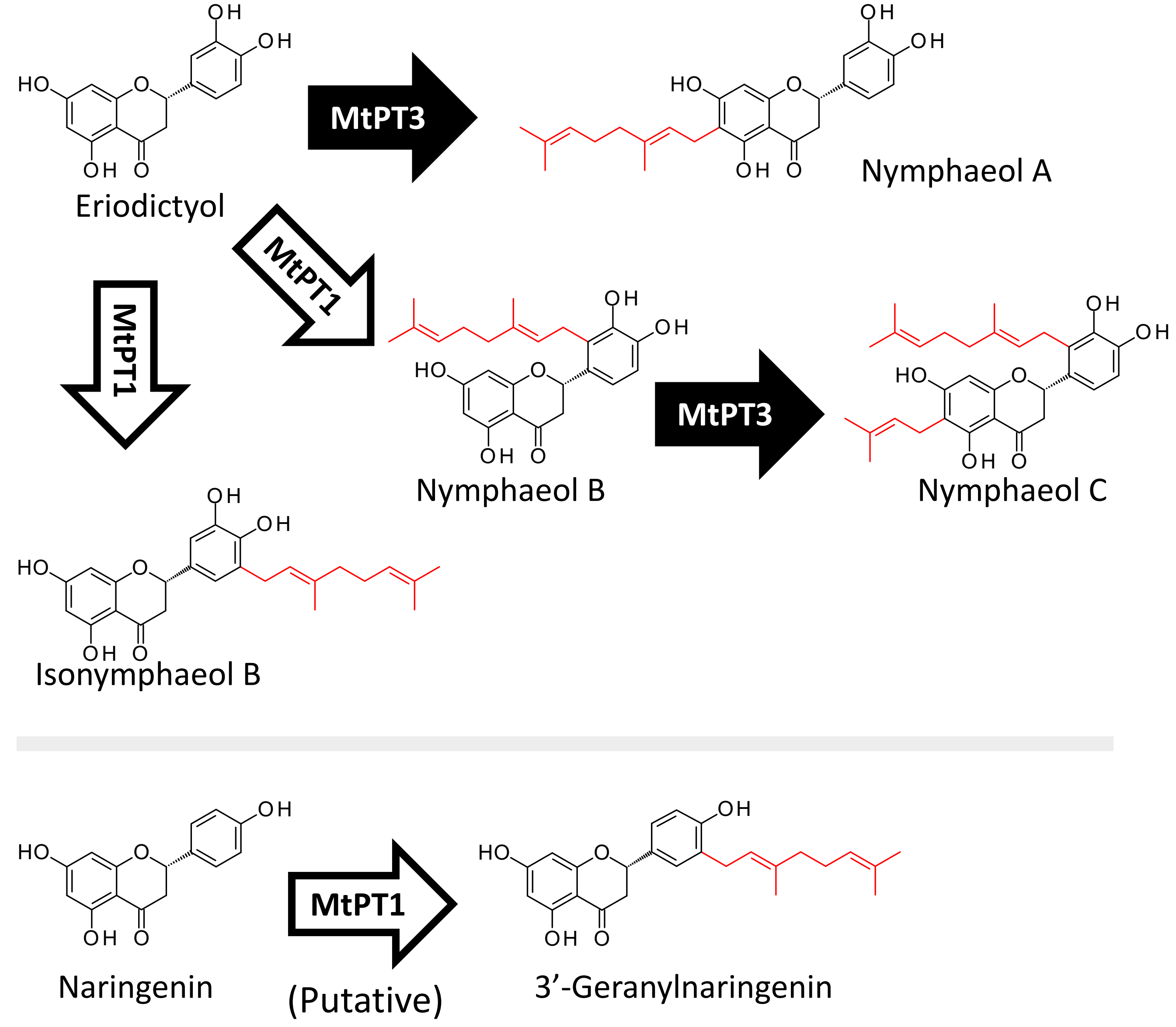

**Supplementary Figure S9. Proposed functions of MtPT1 and MtPT3 in *M. tanarius* glandular trichomes.**
